# Generation of a robust reference gut microbiome dataset for an urban population in Argentina optimized by a machine learning approach

**DOI:** 10.1101/2023.06.24.546376

**Authors:** Cristian Rohr, Mariela Sciara, Bianca Brun, Fabian Fay, Martín P. Vazquez

## Abstract

Robust human microbiome analysis requires robust reference datasets obtained from a population that presents similar habits to the one we are trying to assess.

We reported here the construction of a robust reference dataset of healthy individuals from urban and surrounding rural areas of the Argentine population. We screened 200 volunteers with strict inclusion/exclusion criteria. Volunteers were also screened with routine blood clinical test analysis and a complete metabolome profile from blood and urine to remove outliers before inclusion in the Next Generation Sequencing dataset. Sequencing was done on an Illumina MiSeq using the V3-V4 16S rRNA. Using these data, we performed de novo community structure prediction by applying clustering methodology based on seven distance and dissimilarity metrics and two clustering methods to the reference set. Using this approach, we discovered four different enterotypes in this community structure. We then trained a model for the classification of any new sample into the structure of the reference set. Once the new sample was classified, it was compared to the reference ranges of both the enterotype-specific subset and the whole reference set.

Finally, we challenged the robustness of this methodology using samples from two test case volunteers with clinically proven gut dysbiosis in a time-series sampling with dietary interventions. Our results pointed to the need to carefully analyze the results of gut microbiome in the context of enterotype-specific rather than to a whole population dataset.

## 1. Introduction

The Microorganisms in the gastrointestinal (GI) tract play a role in nutrient uptake, vitamin synthesis, energy harvest, inflammatory modulation, and host immune response, among other functions. Important factors such as age, birth delivery route, antibiotic use, and diet can shape the gut microbiota and they are even more important than host genetics in doing so (1, 2, 3)

Reproducible patterns of variation in the microbiota have been observed in the adult human gut (4). When separated into clusters, these variations were termed enterotypes and proposed as a useful method to stratify human gut microbiomes. However, due to the nature of microbiome clustering in the gut, the number or even existence of different community types has been a topic of heated debate after the publication of the original work (4). An analysis of the three largest publicly available datasets and reports from different publications, consistently reveals that the local substructure is always similar, that is a three-cluster model finds Bacteroides, Prevotella and Firmicutes dominated clusters.

Holmes et al. (5) propose an alternate approach to identify structure. Their method identifies a generative model for each possible state and determines how each one explains the observed data, focusing on the actual genera abundances rather than the distances. Using this approach (Dirichlet multinomial mixture models; DMMs) they reported four generative processes. Two of the clusters consisted of Bacteroides and Prevotella enterotypes, while a third showed an increased prevalence of Ruminococcus and other Firmicutes genera. The last cluster had a high fraction of unidentified taxa. A further study that applied the same method to the HMP 16S rRNA data found that the gut microbiome is best approximated with four similar models. Two of these are overlapping with Bacteroides and Prevotella enterotypes, while the other two are a more complex mixture.

Thus, independent of clustering and modelling approaches, bacterial co-abundance networks provide a species network that may underline the fundamental properties of these preferred community profiles.

Therefore, there is certain agreement that there are distinct areas within the complex microbial composition landscape in which the respective gut communities show biological differences. The concept of enterotypes can help capture such differences, although defining meaningful and robust boundaries remains a challenge (4).

Bacteroides and Prevotella are two well-known enterotypes in human gut microbiomes, with Fimicutes (ruminococcus, faecalibacterium or others) being a third, and a complex fourth cluster or enterotype is also recognised. In any case scenario, There is always a particular enterotype dominating over the others but they may dynamically change according to specific environmental drivers.

As mentioned, diet and environment dominates over host genetics in shaping the human gut microbiome (3,6,7). In consequence, a reference dataset should be constructed for a fine-tuning analysis of every different geographical region and/or lifestyle.

With that goal in mind, we published a pilot study from 20 healthy volunteers of the city Rosario in Argentina in 2016 (8). Results obtained showed that Argentinean microbiotas were different from the microbiotas of HMP in community composition. This pilot study reinforces the importance of analyzing an adult cohort to define a reference dataset for microbiome analysis in different populations.

In the present work, we reported the construction of a robust reference dataset of healthy individuals for urban and surrounding rural areas of the Argentine population. We applied machine learning models for clustering and classification and discovered four different enterotypes in the community structure. Finally, we challenged the robustness of the reference set against two test case reports.

## 2. Materials and methods

### 2.1. Control volunteers recruitment and study design

The target was to recruit 200 healthy volunteers for sampling as a reference dataset for gut microbiome analyses. Volunteers were recruited from four different rural areas and urban cities of Argentina as follows:

| City | Population | localization | Area |
| --- | --- | --- | --- |
| Rosario | 1,240,000 | 32.9587° S,<br>60.6930° W | 180 Km2 |
| Rafaela | 106,000 | 31.2526° S,<br>61.4916° W | 156 Km2 |
| Paraná | 248,000 | 31.7413° S,<br>60.5115° W | 137 Km2 |
| Venado Tuerto | 76,000 | 33.7456° S,<br>61.9690° W | 47 km2 |

The recruitment of healthy volunteers was conducted as an open citizen science project with awareness in the public media and social media networks. The project was open from December 2015 to december 2016. Volunteers were instructed about the project and directed to any of the four clinical labs in the different cities participating in the project for sampling and signing the informed consent agreement. Inclusion and exclusion criterias were as follows:

#### Inclusion criteria

1. Adult, healthy, male or female, between 18 and 50 years of age.
2. Physical examination with no findings of clinical relevance
3. NON-REACTIVE serology for: HBV surface antigen (HBsAg), antibodies against human immunodeficiency virus (anti-HIV) and HCV antibodies (anti-HCV).
4. In female volunteers, negative blood pregnancy test.
5. Being capable of reading, understanding and signing an Informed Consent. Exclusion criteria
6. Suffering gastrointestinal disorders such as:
7. Chronic intestinal inflammation, ulcerative colitis, Crohn’s disease, or indeterminate colitis; Irritable colon syndrome (moderate to severe); Infectious gastroenteritis, colitis or gastritis; chronic or persistent diarrhoea of unknown origin, recurrent infection with Clostridium difficile or untreated Helicobacter pylori infection; Chronic constipation.
8. Having consumed in the last 6 months prior to the sampling:
  - Any type of antibiotics (oral or intravenous).
  - Any type of corticoid (oral, intravenous, intramuscular, nasal or inhaled).
  - Cytokines or drugs that stimulate the immune system (such as interleukin).
  - Cytotoxic agents (chemotherapy)
  - Probiotics consumed in large doses (tablets or powders). Common dietary components such as yogurts do not apply.
9. Being diagnosed with any disease in the last six months.
10. Chronic use of any type of medication.
11. Have a vegan or vegetarian diet as a habit.
12. Have a Body Mass Index (BMI) >29.99 - obese patients.

### 2.2. Biological Sample collections from volunteers

200 volunteers were screened during the timeframe where the project was open. Several individuals were dropped at screening phase for not meeting strict inclusion/exclusion criteria. The project ended up with 172 volunteers to proceed with biochemical and metabolome analyses from blood and urine samples, and microbiome analyses from stool samples. These represent a total of 516 biological samples to process in the laboratory.

Epidemiological and contextual metadata were collected for each volunteer using a questionnaire (see supplementary file 1) for general background information, diet, clinical history, gestational history, and medication history

Blood samples and urine samples were collected at clinical lab locations in every city from each participant to determine clinical biochemistry and metabolome data. The goal of these data was to identify other potential outliers, besides inclusion/exclusion criteria, that needed to be excluded from the final reference set.

Stool samples were collected at home using a swab with a provided kit. Colector tubes containing the swabs were conserved in a zymo buffer at 4C in the home refrigerator until collected by clinical lab personnel and immediately processed for DNA extraction.

### 2.3. Biochemical and Metabolome analysis

Blood, urine and stool samples from all volunteers were sent to the reference lab in Rosario city (CIBIC laboratories, Rosario, Argentina. www.cibc.com.ar) for initial processing and biobank storage.

Blood and urine samples were processed for clinical biochemistry analysis in CIBIC labs. A total of 21 clinical determinations were performed per sample as detailed in supplementary file 2.

Plasma and urine samples were randomized and sent for metabolome analysis to PLABEM (Plataforma Argentina de Biología Estructural y Metabolómica, Rosario, Argentina), (www.plabem.gob.ar). In brief, 300 ul samples were mixed with equal amounts of buffer and transferred to NMR tubes. All samples were processed for acquisition of CPMG, NOESY and JRES NMR spectra with phase correction, baseline correction, normalization, non-informative regions elimination and extraction of spectral data. NMR spectra were analyzed in the context of the available metadata collected from the volunteers (see supplementary file 1)

### 2.4. DNA extraction and DNA Sequencing

Total DNA extraction from stool samples (about 200 mg) was performed in triplicates using QIAmp DNA Stool Minikit following manufacturer’s instructions. The 16S rRNA V3-V4 hypervariable region was first amplified using PCR method (20 cycles) and then a second round for sample identification (6 cycles) was performed. Amplicons were cleaned using Ampure DNA capture beads (Argencourt-Beckman Coulter, Inc.) and quantified using Quanti-iTTM PicoGreen® DNA Assay Kit (Invitrogen Molecular Probes, Inc., Eugene, OR, United States) with the standard protocol (high range curve – half area plate) and pooled in similar concentrations before sequencing on the Illumina MiSeq platform (Illumina, Inc., San Diego, CA, United States) using 2 × 300 cycles PE v3 chemistry. For each sample, three replicates (three extraction from the same biological sample) were made to diminish the effect of amplification bias.

Stool samples from the test cases were processed the same way.

### 2.5. Next Generation sequencing bioinformatics analysis

Amplicon sequencing produced 50 million raw paired-end (PE) reads for a 15 gb of raw dataset. The dataset contained more than 110,000 reads of raw coverage per sample triplicate. Samples were demultiplexed and duplicated reads eliminated. We followed the 16S SOP described in Microbiome Helper (9). FastQC (v0.11.5) was used to analyze the raw data quality of PE reads. Paired-end reads were stitched together using PEAR (v0.9.10). Stitched reads were filtered by quality and length, using a quality score cut-off of Q30 (phred quality score) over 90% of bases and a minimum length of 400 bp. Concatenated and filtered fastq sequences were converted to fasta format and we removed sequences that contained “N”. Potential chimeras were identified using the UCHIME algorithm, and then the chimeric sequences were removed.

Data cleaning yielded 45,000 reads per sample triplicate of effective coverage for a high quality dataset.

The sequences were clustered to operational taxonomic units (OTUs) at 97% similarity level with open reference strategy implemented in Quantitative Insights into Microbial Ecology (QIIME; v1.91), using SortMeRNA for the reference picking against the Greengenes v13 8 97% OTU representative sequences database and SUMACLUST for de novo OTU picking.

Bioinformatics analysis of the data was done using a custom QIIME pipeline and the R package phyloseq (https://github.com/joey711/phyloseq). OTU table was rarefied at 10,000 sequences per sample for alpha and beta diversity and random forest analysis. To provide alpha diversity metrics, we calculated observed species, Chao1 and Shannon’s diversity index. To evaluate beta diversity among cases, controls and healthy individuals UniFrac (weighted and unweighted) distances and Bray Curtis dissimilarity were used, prior removal of OTU’s not present in at least 5% of samples. The UniFrac and Bray Curtis measures were represented by two dimensional principal coordinates analysis (PCoA) plots. Differences between groups were tested by a permutation multi-variate analysis of variance (PERMANOVA) using distance matrices function (ADONIS) implemented in the R vegan package [10, 11]. The evaluation of beta diversity between cases and controls were also determined using principal fitted components for dimension reduction in regression as it was used to analyze microbiome data previously [10,11].

There was a very good correlation among triplicates. In the end, triplicates were pooled together and the average value used for the dataset.

Test cases were processed in the same exact way and compared to the reference set

### 2.6. Machine learning optimization

*Preprocessing for machine learning*. OTUs were normalized using Total Sum Scaling normalization (TSS) after rarefaction. A taxonomic filtering was applied, OTUs not classified at genus level were removed, and remaining ones were grouped (different OTUs that belong to the same genus were joined).

#### Unsupervised learning (Clustering)

We followed a methodology as proposed previously [12,13]. Two clustering algorithms PAM [12, 13] and hclust were applied to the preprocessed OTU tables. For both clustering algorithms, we evaluated seven distance/dissimilarity metrics: Jensen Shannon Divergence and his root square version (JSD; rJSD), Bray Curtis, Morisita-Horn, Kulcynsky, UniFrac and weighted UniFrac [10]. To assess the quality of the clusters, the coefficient of Silhouette [14] and Prediction Strength [15] scores were used followed by an additional bootstrapping step assessed with Jaccard’s similarity score [16]. In each case, possible values of the number of clusters k from 2 to 10 were evaluated. The proposed methodology begins by searching for clusters with the two algorithms (PAM and Hclust), with the different distance metrics, and all the possible values of k. In total, 126 different clusters were compared (2 algorithms, 7 distance measurements, 9 k values). To select the best result we used the following criteria:

1. Use k clusters (from 2 to 10) with the highest average Silhouette width coefficient with the score above 0.25
2. Check if the value of k selected above has a Prediction Strength value greater than 0.8, or
3. if the cluster is stable according to Jaccard, use a value of 0.75 in a bootstrapping process.

##### Supervised learning

Once the clusters were obtained, they were used as class labels for each individual to train supervised learning models. These models used genus level abundance as features to learn how to assign a given sample to a group. The objective of these models is to identify the group of an unknown sample, and compare it with this specific group. For that purpose, we implemented random forests and support vector machines algorithms.

The dataset of control individuals (n = 152) were partitioned in 70% training and 30% testing. The model evaluation was performed using a 10 fold cross validation repeated 3 times to select for the best parameters in the training dataset and the validation dataset to obtain the best possible model. This was done using the caret 6.0-81 library in R (http://topepo.github.io/caret/index.html).

##### Oversampling

To address the imbalance of classes in the training data set, we used an over-sampling algorithm called ADASYN (Adaptive Synthetic Sampling Approach for Imbalanced Learning) from the R smotefamily 1.3 library (https://cran.r-project.org/web/packages/smotefamily/index.html). This algorithm generates new instances for the minority class (s), instead of replicating instances of oversampling. *Hyperparameters optimization* - For RF and SVM algorithms, different combinations of parameters were evaluated and selected as the best combination considering the results on the training dataset after performing cross validation on 10 partitions repeated 3 times. In the case of Random Forests, the parameter mtry (number of variables randomly taken as candidates in each split) was tuned according to the sequence from 1 to the total number of features with a step of 7 and the parameter ntree (number of trees) was set with values 500, 1000, 2000, 5000. In the case of SVM, the radial, poly and linear kernel type parameters were set to cost (0.5, 1, 2, 4) and gamma (4, 8, 16).

### 2.7. Case studies

Participants LB and FG volunteered in spontaneous presentation to the open science project to analyze their gut microbiomes due to concurrent gastrointestinal symptoms. They signed the informed consent agreement as the rest of the healthy volunteers. In the case of LB, three samples were taken at days 1, 7 and 14. In the case of FG, three samples were taken on days 1, 15 and 30. In both cases, each sample was extracted and processed in triplicate.

## 3. Results

### 3.1. Construction of the Reference set sample for gut microbiome

A total of 200 volunteers were screened from four different cities of Argentina in an open science project to collect blood, urine and fecal samples to build a robust reference set for gut microbiome analyses (Fig. 1A).

**Figure 1.**
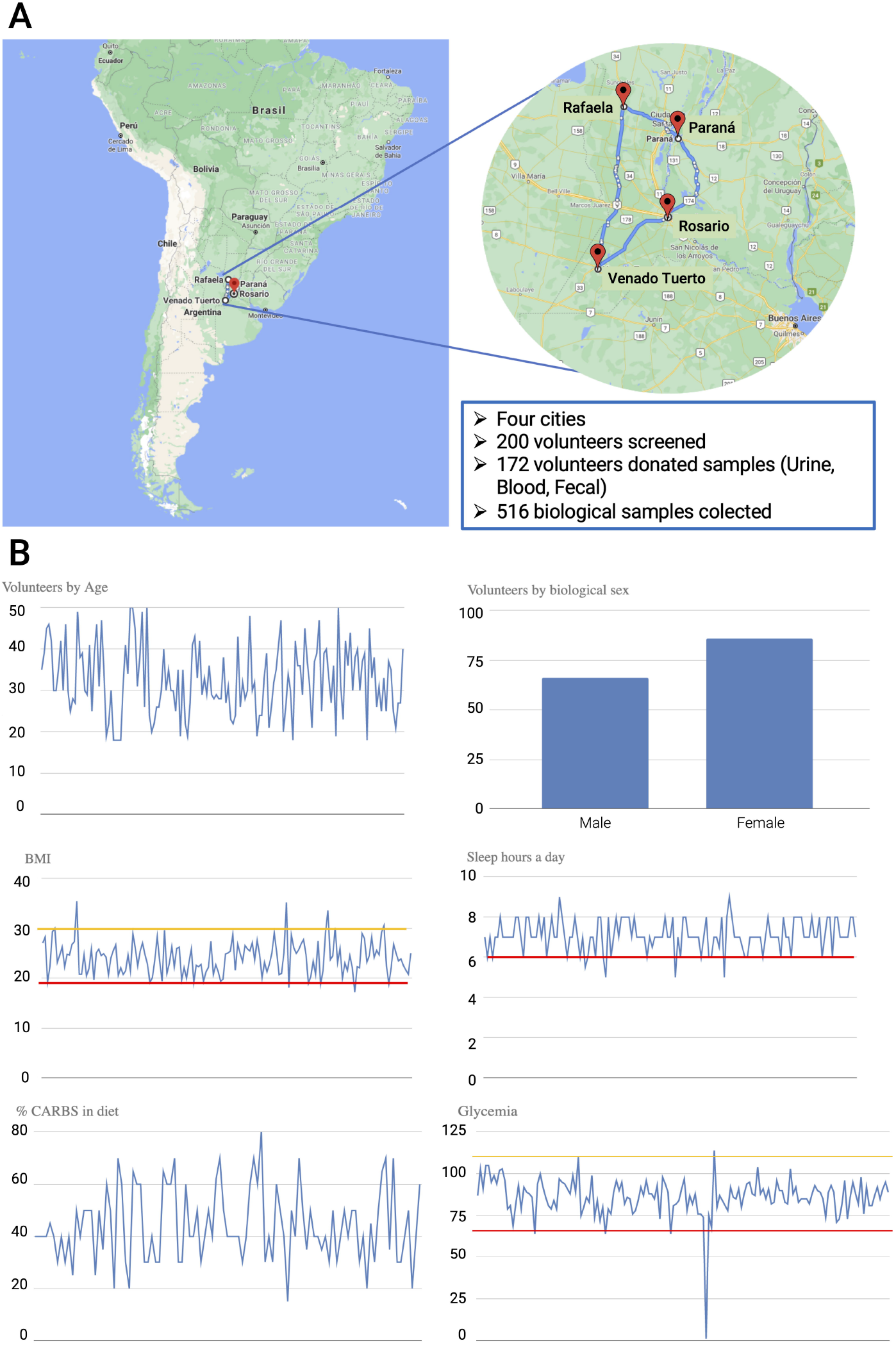
A. Map details showing the location of the cities that participated in the project. B. Graphical representation of all volunteers organized by age, biological sex, body mass index (BMI), sleep hours per day, amount of declared carbohydrates in diet daily, and glycemia determined in the clinical lab. Yellow lines represent upper threshold limits and red lines lower threshold limits

172 volunteers passed the initial screening process (18 to 50 years old). Then, three biological samples (blood, urine and fecal) were collected per individual for a total of 516 unique samples. Volunteers also answered an extensive questionnaire of habits and diet (contextual metadata, S1 file).

Volunteers were classified by age, biological sex, body mass index (BMI), sleep hours per day and percent of manifested carbohidrates in the diet along with their reespective glycemias at the time of sampling (Fig. 1B).

Blood samples were analyzed in routine clinical tests along with the contextual metadata to eliminate outliers with abnormal parameters.

Clinical tests performed included glycemia, total cholesterol, LDL, HDL, triglycerides, creatinine, uremia and minerals potassium, chlorides and sodium. Enzymes activities measured included GOT, GPT, GGT and ALP to estimate overall liver health (S2 file). Volunteers that presented two or more deviations in the clinical measures were flagged as outliers and those with only one deviation were further analyzed in the light of contextual metadata and metabolome data prior to be dismissed as outliers.

Plasma and urine samples were analyzed by metabolome data using NMR CPMG spectra for plasma and NMR NOESY spectra for urine.

Metabolome data were classified into outliers and strong outliers. Strong outliers were immediately removed from the dataset (S3 Fig) and outliers were further analyzed in the context of clinical test results and metadata. Plasma outliers showed correlation with VLDL particles (S4 Fig) and, indeed, identified individuals that presented higher levels of triglycerides in clinical tests. Urine metabolome identified a strong outlier that matched high levels of ethanol and was immediately removed from the set (S5 Fig). Two outliers matched elevated levels of Hippuric acid and another four outliers matched elevated levels of 3-(3-Hydroxyphenyl)-3-hydroxypropanoic acid, a molecule that is usually produced by clostridium sp. in the gut (S6 Fig). To remove outliers, we inspected if the individual matched another deviated parameter in the clinical tests.

Taking all the data into account, 24 outliers were removed. Thus, the final reference dataset comprised 152 volunteer high quality fecal samples flagged for NGS processing.

Samples were sequenced in an Illumina MiSeq in 2×300 PE configuration to produce 50 million reads accounting for 45,000 reads per sample after data cleaning for a high quality dataset.

### 3.2. Bioinformatics analysis of the reference set samples

The sequences were clustered into operational taxonomic units (OTUs) at 97% similarity level with an Open Reference Strategy implemented in Quantitative Insights into Microbial Ecology (QIIME; v1.91) (17). Procedures are described in materials and methods section.

There was a very good correlation among replicates. They were pooled together and the average value was used for the final dataset.

Samples were analyzed using UniFrac and Bray Curtis measures and they were represented by two dimensional principal coordinates analysis (PCoA) plots (Fig. 2) Results indicated a partial clustering of samples into two groups in both unweighted and weighted unifrac methods. Interestingly, Rosario city tends to cluster separately from Rafaela and Venado tuerto while Parana was scattered all over the plot. This pattern explained 28.14% of the variation observed in the PC1 axis of the weighted unifrac, while was only 4.89% in the PC2 axis in the unweighted unifrac. These results suggested that the proportion of OTUs rather than different OTUs was the main component in the observed clustering. Bray-Curtis analysis rendered similar results (S7 Fig). It is worth noting that Rosario samples from this work clustered together with our previous published work from a pilot analysis of 20 samples from the same city (Fig. 2) (8).

**Figure 2.**
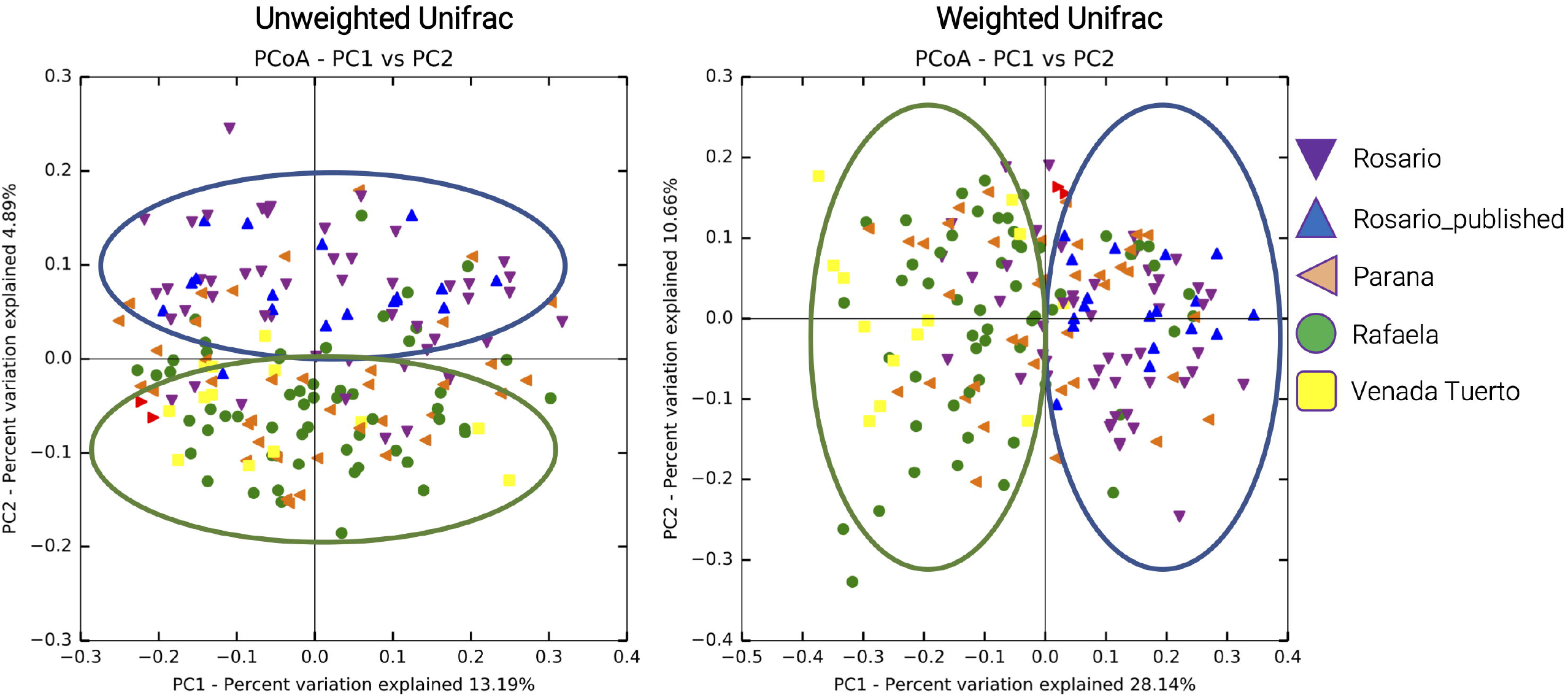
Principal components analysis (PCoA) using unweighted and weighted unifrac of the gut microbiome data obtained from the four cities. Blue diamond labeled Rosario_published corresponds to the gut microbiome from Rosario city previously published by our group using the same methodology (8).

**Figure 3.**
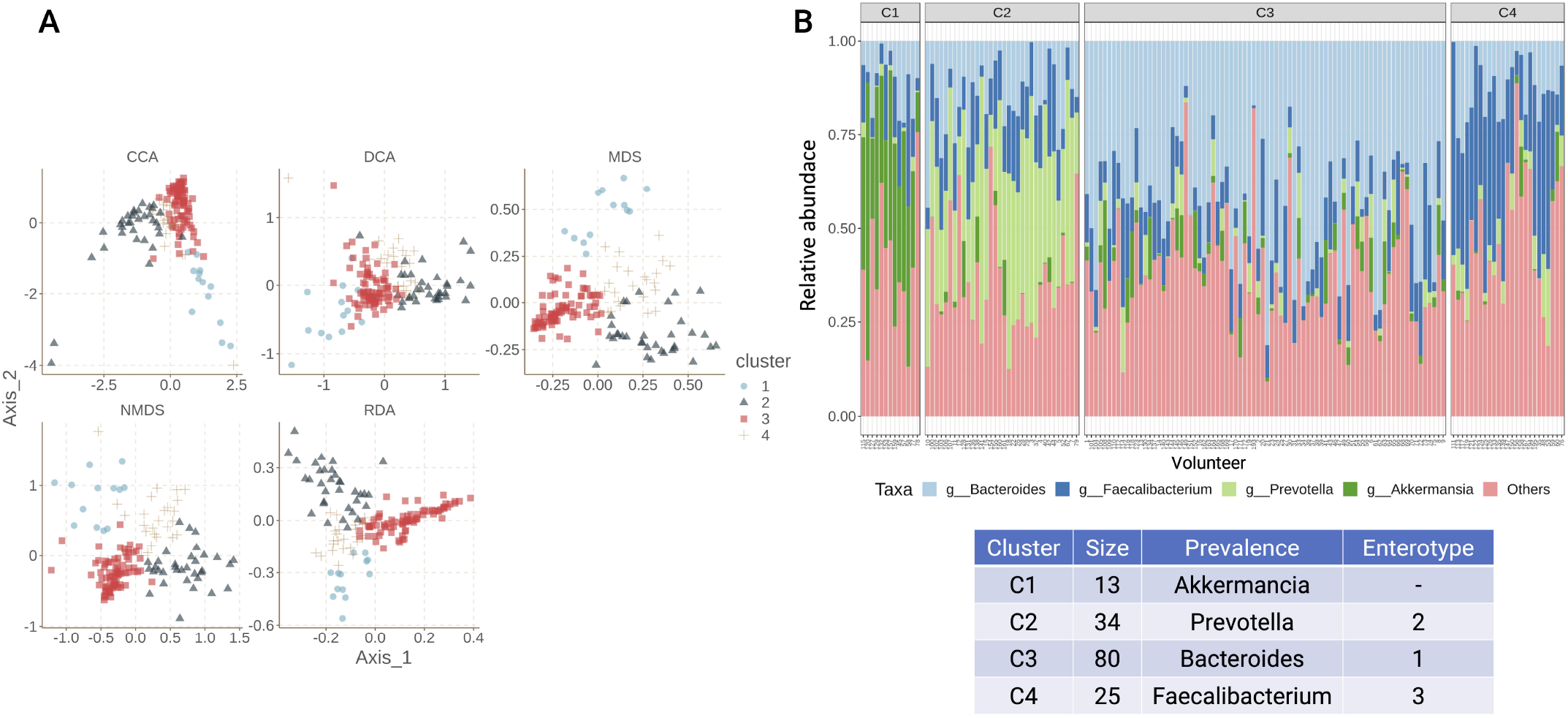
A. Two-dimensional Clustering analysis of the gut microbiome data. CCA: canonical correspondence analysis, DCA: detrended correspondence analysis, MDS: multidimensional scaling, NMDS: non-metric multidimensional scaling,, RDA:Redundancy analysis. B. Relative abundance of the most frequent genera in each cluster. We labeled our de novo clustering discovery as C1, C2, C3, C4. Each column represents a volunteer in the analysis. The table below depicts the prevalence and the representative genera of each cluster as well as the mapping to the enterotypes published (4)

### 3.3. Machine learning optimization of the reference set samples

We investigated the clustering in this reference set since enterotypes were well described in several studies (4).

We applied two clustering approaches, hierarchical and partitional (Hclust and PAM algorithms), to reveal the different composition and variation of the gut microbiome in the reference set. In each case, we considered 9 possible values of partitions from k=2..10 and 7 distance/dissimilarity metrics (JSD, rJSD, BrayCurtis, Kulczynski, MorisitaHorn, UniFrac and Weighted Unifrac). The Morisita Horn distance delivers the best results for both clustering algorithms. Best result was obtained for the PAM algorithm with a value of k=4 partitions. This result was used for the following analysis taking into account the assessment approach proposed in [12]: a) Silhouette coefficient SI value 0.501, b) the prediction strength coefficient did not exceed the cut-off value of 0.8, maintaining a value of 0.516 (however it’s the highest value, except for k = 2 when considering the PAM algorithm), c) the Jaccard score presented a value of 0.794 for the combination of the PAM algorithm with the Morisita-Horn distance, exceeding the cut-off value of 0.75 (S8 Fig).

In sum, we found four clusters in the reference set (Fig. 4A). Each cluster is characterized by the prevalence of a specific genera of bacteria (Fig 4B): C1) Akkermansia, C2) Prevotella, C3) Bacteroides and C4) Faecalibacterium. Our results were in line with previous reports of enterotypes, particularly for enterotype 1 (Bacteroides), enterotype 2 (Prevotella) and Enterotype 3 (Ruminoccoccus). Some reports indicated the presences of a fourth undefined enterotype. In our case, it was defined by the Akkermansia prevalence (Fig 4B).

**Figure 4.**
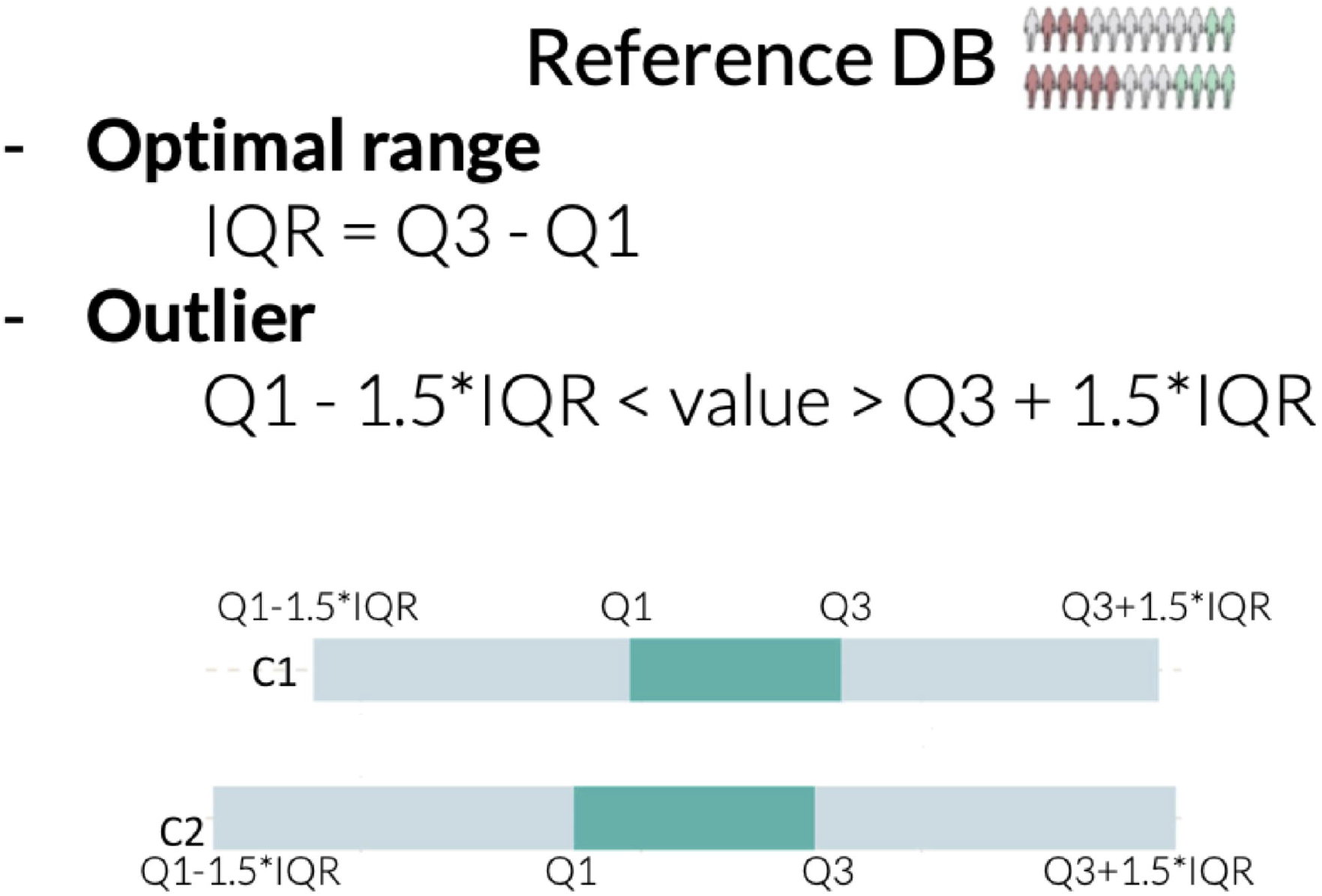
Definitions of the optimal ranges and outliers in our gut microbiome database. IQR: Inter Quartil Range is defined as the values between Quartil 1 (Q1) and Quartil 3 (Q3). Outliers are defined as the values outside de IQR up to a value of 1.5*IQR. Below is the schematic representation of the normal range and outliers for cluster specific databases C1 and C2 as an example.

We trained Random Forests and Support Vector Machines algorithms on reference samples. The purpose of these models was to classify any given new sample from individuals of similar populations into the observed clustering structure of the reference set.

The best model obtained was using the Random Forests algorithm with oversampling with an Accuracy of 0.977 and scores of 0.9885 and 0.9519 for micro and macro F1 respectively. Using oversampling allowed us to obtain better metrics for both algorithms. The results of the parameter optimization process on the set of training data with oversampling for the Random Forests algorithm are shown in Fig. S9. The relevance of the different genera to classify the samples evaluated according to the GINI index are shown in Fig. S9. Indeed, the genera that we found to characterize the four clusters are the ones that were more relevant to classify the samples.

Finally, we defined the reference range values using the reference set samples for a variety of microbiome markers such as shannon diversity index, F/B ratio, Phylum, family and genus proportions. This was a very important definition in order to compare any new sample against the reference set. To that end, we defined reference range values for the complete set (all DB) and for each one of the four clusters individually. The optimal range values were defined as the interquartile range IQR=Q3-Q1 and denoted by a green box limited by the marks Q1 and Q3. The outlier range values were defined as Q1-(1.5*IQR)<Value>Q3+(1.5*IQR) and denoted as light green boxes surrounding the optimal range (Fig. 4). Any new sample value will be marked by a black dot in this type of graphics to easily visualize if it is in the range or out of the range compared to the reference set samples. One important issue to address is if the value of a new sample needs to be interpreted when compared against all DB or against its specific cluster DB. To investigate this, we evaluated test case volunteers.

### 3.4. Test cases studies

Test case volunteers allowed us to challenge the quality of the microbiome reference dataset against clinically diagnosed individuals with symptoms compatible with gut dysbiosis.

We allowed two volunteers, LB and FG, into the citizen science project that consented to participate.

Volunteer LB was a 33-years old female living in Rosario city with at least five years of documented gastrointestinal (GI) disorders. She presented persistent abdominal pain and bloating with occasional diarrhea and vomiting. She also manifested anxiety and depression episodes related to her clinical GI symptoms. There were no relevant altered parameters in the clinical lab testing at the time of sampling. However, she was on medications such as 40mg of Pantoprazole to treat reflux symptoms and 120 mg Simeticone + 200 mg Trimebutine to treat abdominal pain. The diet was primarily vegetarian with occasional inclusion of chicken and fish two times a week. This diet was present for at least five years prior to this project.

LB donated three fecal samples to the project at seven days intervals (day 1, day 7, day 14). During the sampling, she manifested the GI symptoms of abdominal pain and bloating. Immediately after day 1 sampling, she followed a strict diet of gluten-free, lactose-free meals composed mainly of Arugula, carrot, pumpkin, cabbage, yamani rice and abundant liquids. Samples were processed in the lab and sequenced as described above in the same conditions as those of the reference set.

The LB samples were clearly outgroups when compared to the reference set in PCoA plots using Bray-Curtis or Unifrac distances (S10 Fig)

Shannon diversity index showed consistent results across the three samples in 14 days (4.42, 4.58, 4.26 respectively) and they were normal values in the upper quartile in the reference set, suggesting that LB microbiome presented a good diversity in the gut microbiome.

Day 1 sample belonged to our C4 cluster (Faecalibacterium, Ruminoccoccacea), while day 7 sample to the C3 cluster (Bacteroides) and finally day 14 sample to the C2 cluster (Prevotella).

However, the F/B ratio in day 1 sample (value 3.79) was out of range when compared to the complete reference set but not if it was compared to the cluster of belonging. In contrast, the F/B ratio in day 7 sample (value 2.59) was almost normal considering the complete reference set while was elevated considering only the cluster of belonging. Last, F/B ratio in day 14 sample (value 1.85) was in the normal range compared to the complete reference set but it was elevated considering only the cluster of belonging (Fig. 5A).

**Figure 5.**
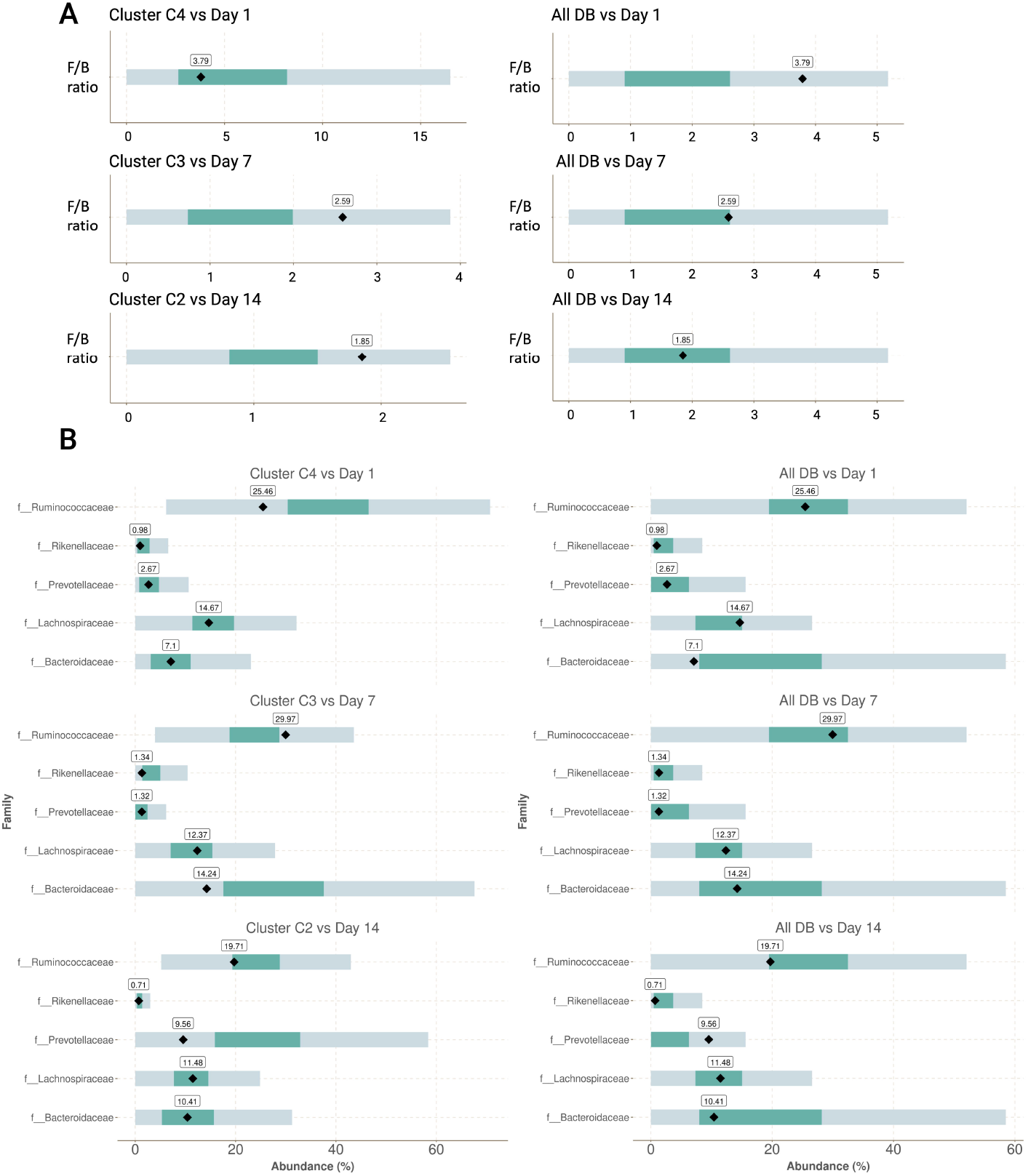
Test case analysis of gut microbiome (LB participant). A. Firmicutes/Bacteroidetes ratio (F/B ratio) at days 1, 7 and 15. Left side, comparison against cluster-specific database. Right side, comparison against the complete database (All DB). B. Family-level analysis of the gut microbiome of LB volunteer at days 1, 7 and 15. Left side, comparison against cluster-specific database. Right side, comparison against the complete database (All DB). The value obtained from the volunteer in each case is indicated by a black diamond.

**Figure 6.**
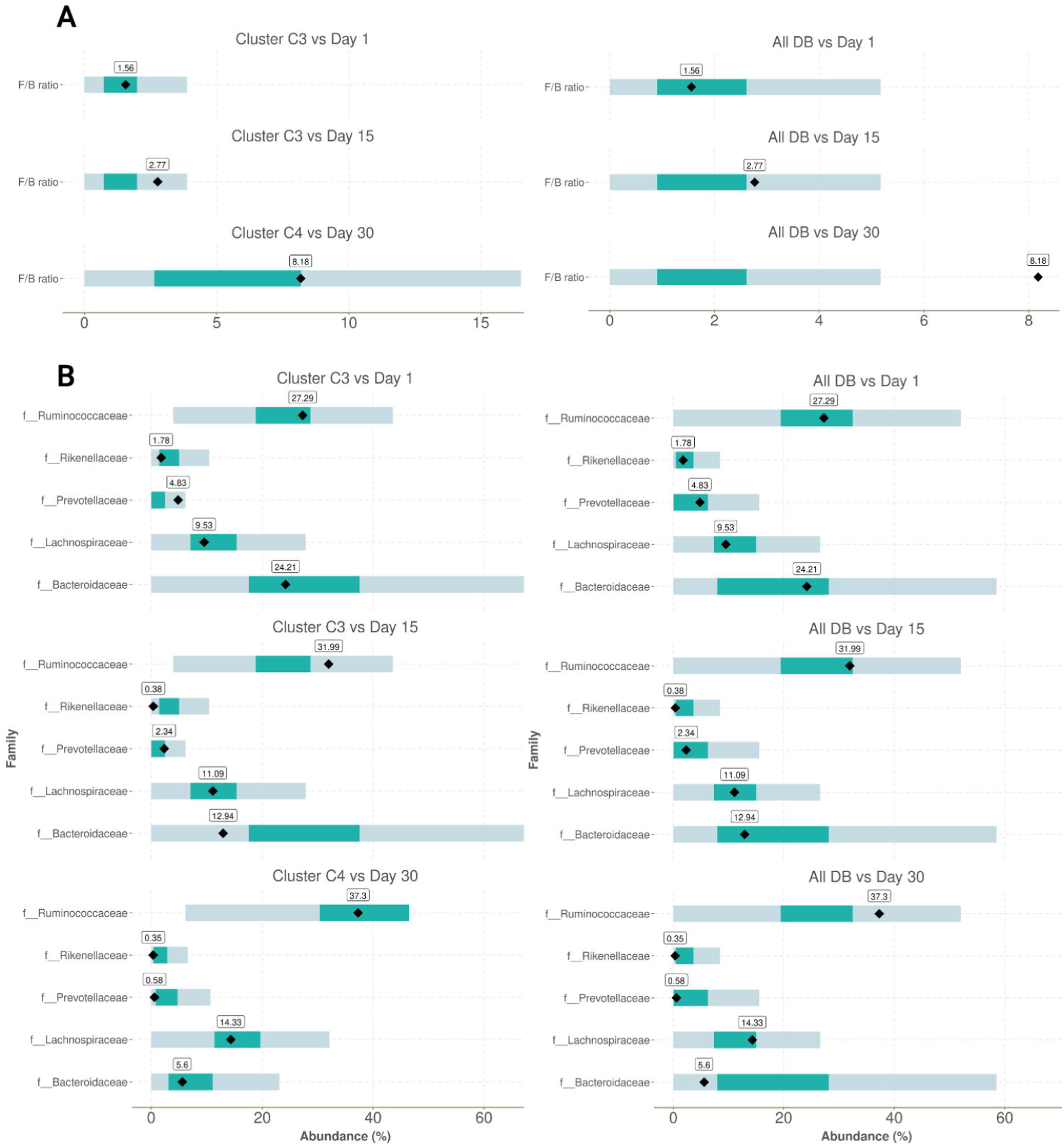
Test case analysis of gut microbiome (FG participant). A. Firmicutes/Bacteroidetes ratio (F/B ratio) at days 1, 15 and 30. Left side, comparison against cluster-specific database. Right side, comparison against the complete database (All DB). B. Family-level analysis of the gut microbiome of FG volunteer at days 1, 15 and 30. Left side, comparison against cluster-specific database. Right side, comparison against the complete database (All DB). The value obtained from the volunteer in each case is indicated by a black diamond.

In this way, one must be cautious interpreting F/B ratios in the light of these results and considering the reference set of comparison (All DB vs cluster DB) (Fig. 5) Nevertheless, the absolute values for F/B ratio in the three consecutive samples of LB in the 14 days period was consistently decreased after following the strict diet (values 3.79, 2.59 and 1.85, respectively)

A detailed inspection at the family level showed that this decrease in the F/B index was mainly correlated with an increase in the levels of Bacteroideacea at day 7 (values 7.1 and 14.24, respectively for day 1 and day 7) followed by an increase in Prevotellaceae at day 14 (values 1.32 and 9.56 respectively for day 7 and day 14) (Fig 5B). This result was in line with the type of diet followed by the participant during the 14 days period.

Volunteer FG was a 33-year old male resident in a rural area of the province of Buenos Aires. He was a lean participant with a BMI of 19. FG was a high performance athlete throughout highschool until 20 years-old, then dropped but continued with moderate physical activity three times per week. After dropping high performance, problems began in the following years such as chronic fatigue, anxiety disorders and bad dietary habits. Gastrointestinal symptoms began and continued to the present day worsening year over year with persistent flatulence, burping, bloating, and alternating constipation and diarrhoea. In the 2-years prior to participating in this study, he was diagnosed with a Candida infection and he followed a strict diet since then. Dietary habits are mainly vegetarian including lettuce, fennel, carrots and whole grains. Dietary restrictions included gluten, lactose and refined carbohydrates.

FG donated three fecal samples to the project taken in 15 day intervals (day 1, day 15 and day 30). During the sampling, he manifested some abdominal discomfort such as pain and bloating and he followed a strict month diet with increasing amounts of whole grains and fibers in the meals.

Samples were processed in the lab and sequenced as described above in the same conditions as those of the reference set.

The FG samples were outgroups when compared to the reference set in PCoA plots using Bray-Curtis or Unifrac distances.

Shannon diversity index was consistent across all samples with values 4.52, 4.36 and4.33 respectively located in the upper third quartile when compared to the complete reference set. FG samples showed no apparent issues at the diversity levels despite his prior history of candida infection (Fig. 7A)

Day 1 and day 15 samples were determined as cluster 3 (Bacteroides), while day 30 sample was determined as cluster 4 (Faecalibacterium, Ruminoccoccacea).

The F/B ratio value was increased successively during the sampling period from 1.56 to 2.77 to 8.18 respectively. The value was in the normal range for day 1 sample either compared to complete reference set (all DB) or to cluster-specific subset. In contrast, day 15 sample was an elevated value in either comparison. Finally, day 30 sample value was almost in the normal range compared to cluster-specific subset but was way out of range when compared to all DB. Once again, these results call to attention the need to carefully evaluate the values of F/B ratio in perspective of the comparison dataset (Fig. 7A).

When inspected at the family level, it was observed a successive increase in the values of Ruminococcaceae (27.56, 31.79, 37.11) and Lachnospiraceae (9.34, 11.03, 14.2). These families belonging to the phylum Firmicutes were probably the main forces behind the dynamics of migration from cluster 3 to cluster 4. However, it was also noted that prevotellaceae and bacteroidaceae were decreased by 14% and 23.6% respectively and probably contributed to the migration (Fig. 7B)

These results were in line with the dietary habits during the period of sampling that incorporated incremental amounts of whole grains and fiber.

## 4. Discussion

In this study, we presented a complete reference set of samples for gut microbiome analysis based on volunteers from cities and surrounding rural areas in Argentina. One of the main strengths of this reference dataset is that it is highly curated in terms of the volunteers that donated the samples. This specific dataset was constructed using the V3-V4 regions of the 16sRNA gene biomarker for bacteria and it does not include archaea or eukarya domains. However, DNA from all curated samples was stored in a biobank at -70C and it will be used to construct another reference dataset using a metagenome shotgun shallow sequence to include all three domains of life in the near term.

The samples in the dataset were curated taking into account clinical lab testing from blood samples, metabolome testing from plasma and urine samples and contextual metadata from all the volunteers screened to exclude outlier samples. In this way, we kept only the samples that best represented what could be considered a gut in “eubiosis-state” representative of western populations in big cities and surrounding areas.

We stress the importance of curating a high quality dataset and the remarkable need to eliminate potential outliers because of the high variability of the gut microbiome due to environmental factors.

It should also be stressed that our reference dataset is intended to be used in populations similar to the one described here: western industrialized cities and their surrounding rural areas with a major component of european descent. It is clear that other populations will differ from this reference set as their habits and environments differ (1, 3).

It is interesting to note that one of the clusters described in our work were characterized by a prevalent abundance of Akkermansia genus. This genus was dubbed as one of the new microbial populations fuelled by the western industrialized lifestyle (18).

Indeed, western microbiome’s populations are consistent with the relative paucity of dietary fibre consumed. In these communities, an elevated proportion of mucus-degrading enzymes and mucus-consuming species, such as Akkermansia, is observed (18). The other three clusters characterized in this work were consistent with previous observations in industrialized populations (4)

It was proposed that this new microbial community structure along with the disappearance of the so-called VANISHED taxa could be responsible for the modern chronic diseases and low-grade chronic inflammation observed in industrial populations. This is in part because the newly adapted microbiome is not fully compatible with our genome evolution to counteract the changes (18).

A recent published work, elegantly showed how industrialization could impact our gut microbiomes by studying the Irish Traveller community, which has recently undergone enforced lifestyle changes (19). Indeed, the shifts in the microbiome in those Travellers who are more rapidly adopting industrialized practices show many features of the gut microbiomes linked to chronic diseases such as obesity and heart disease (19). The authors also observed that housing conditions, pet-ownership, number of siblings, Body mass index and other factors were important in transition microbiomes besides diet.

While these newly adapted microbiomes to western lifestyle could be considered in eubiosis for the current environment, it is certainly not when compared to ancestral populations as VANISHED taxa were lost (18). However, it is very important to determine what is the structure of this new “eubiosis-state” with the most possible accuracy in order to determine what actually is a new “dysbiosis-state” for a specific western population. There is no single globally reference dataset that will suit all comparison purposes for all western populations. There is a need to determine reference datasets por specific subpopulations according to their diets, lifestyles and housing conditions (1, 2).

Our de novo community structure by machine learning and classifier based on random forest algorithms identified 4 clusters as mentioned above. Costea et al. (4) suggested that when de novo community structure was performed, and for the purpose of standardization, the results should be compared with the procedure and classifier that they described in http://enterotypes.org. Our results were similar using both classifiers (this paper and 4): 88.8% of our samples looked similar to stool samples from large scale projects like MetaHIT and HMP. The bacteroides enterotype (C3 cluster) was 53% and 59.8% respectively. The Prevotella enterotype (C2 Cluster) was 22.4% and 31% respectively. The Firmicutes enterotype (C4 cluster) was 16% and 9.2% respectively. Last, the remaining 7% community proportion was Akkermansia (Verrucomicrobia) in our reference set and 11% of samples were not classified by the algorithm of Costea et al (4).

Finally, the clustering and machine learning optimization of our highly curated microbiome reference set was the key to define the normal ranges of abundance of different bacteria and biomarkers. This, in turn, will be useful to compare other individuals that may present deviations in the composition of their microbiomes or dysbiosis-states. The use of a robust clustering methodology allowed us to classify our gut microbiome reference database into four well-defined groups along with their normal reference ranges. These cluster-specific databases showed us the importance of analyzing microbiome results in the context of the individual’s current enterotype, since what may seem out of range compared to a general population may not be so in relation to its current enterotype. This was the situation observed when analyzing our case test studies. For example, the LB volunteer’s microbiome dynamics was moving from cluster C4 to C3 to C2 during the times of sampling, while the F/B ratios were decreasing. However, day 1 F/B ratio was in range considering the cluster-specific DB but it was out of range considering All DB. On the contrary, while the F/B ratio seemed approaching normal range values in day 7 and day 14 against All DB, it was in fact out of range compared against the specific-cluster DBs (Fig xx). On the other hand, the Shannon diversity index was indifferent to the comparison against All DB or Cluster-specific DB. We call to attention that F/B ratios and bacterial abundances must be carefully evaluated in the context cluster-specific DBs instead of All DBs. Other metrics such as diversity indices may not be affected. Indeed, we carefully selected the test cases and may not represent a general situation but still the data point in that direction.

In sum, our results reinforce the importance of adequate and robust control groups based on the geographic location, housing conditions and diet to characterize other individuals with potential “dysbiosis-states” of modern western populations.

## Supporting information

Rohr_et_al_suppl

## 5. Acknowledgements

We thank all the volunteers that participated in the study. We thank the clinical labs in the cities that participated in this open science project. We are grateful to Drs.

Alejandro “Tato” Vila and Rodolfo Rasia from IBR-PLABEM (https://www.plabem.gob.ar) for providing the metabolome data for the project.

## 6. Author contributions

CR and MPV compiled and analyzed all the data. CR developed the machine learning models. MS and BB collected all the data. MS organized the protocol, questionnaires and the relation with clinical labs and samples logistics. FF and MPV designed the project. MPV wrote the manuscript.

## Notes

### Competing Interest Statement

The authors have declared no competing interest.

## References

1. Gupta, V., Paul, S., Dutta, C. (2017). Geography, Ethnicity or Subsistence-Specific Variations in Human Microbiome Composition and Diversity Frontiers in Microbiology 8(), 229 –16. https://dx.doi.org/10.3389/fmicb.2017.01162

2. Senghor, B., Sokhna, C., Ruimy, R., Lagier, J. (2018). Gut microbiota diversity according to dietary habits and geographical provenance Human Microbiome Journal 7–8(), 1-9. https://dx.doi.org/10.1016/j.humic.2018.01.001

3. Rothschild, D., Weissbrod, O., Barkan, E., Kurilshikov, A., Korem, T., Zeevi, D., Costea, P., Godneva, A., Kalka, I., Bar, N., Shilo, S., Lador, D., Vila, A., Zmora, N., Pevsner-Fischer, M., Israeli, D., Kosower, N., Malka, G., Wolf, B., Avnit-Sagi, T., Lotan-Pompan, M., Weinberger, A., Halpern, Z., Carmi, S., Fu, J., Wijmenga, C., Zhernakova, A., Elinav, E., Segal, E. (2018). Environment dominates over host genetics in shaping human gut microbiota Nature 555(7695), 210. https://dx.doi.org/10.1038/nature25973

4. Costea, P., Hildebrand, F., Arumugam, M., Bäckhed, F., Blaser, M., Bushman, F., Vos, W., Ehrlich, S., Fraser, C., Hattori, M., Huttenhower, C., Jeffery, I., Knights, D., Lewis, J., Ley, R., Ochman, H., O’Toole, P., Quince, C., Relman, D., Shanahan, F., Sunagawa, S., Wang, J., Weinstock, G., Wu, G., Zeller, G., Zhao, L., Raes, J., Knight, R., Bork, P. (2018). Enterotypes in the landscape of gut microbial community composition Nature Microbiology 3(1), 8–16. https://dx.doi.org/10.1038/s41564-017-0072-8

5. Holmes, I., Harris, K., Quince, C. (2012). Dirichlet Multinomial Mixtures: Generative Models for Microbial Metagenomics PLoS ONE 7(2), e30126. https://dx.doi.org/10.1371/journal.pone.0030126

6. Rinninella, E., Raoul, P., Cintoni, M., Franceschi, F., Miggiano, G., Gasbarrini, A., Mele, M. (2019). What is the Healthy Gut Microbiota Composition? A Changing Ecosystem across Age, Environment, Diet, and Diseases Microorganisms 7(1), 14. https://dx.doi.org/10.3390/microorganisms7010014

7. David, L., Maurice, C., Carmody, R., Gootenberg, D., Button, J., Wolfe, B., Ling, A., Devlin, A., Varma, Y., Fischbach, M., Biddinger, S., Dutton, R., Turnbaugh, P. (2014). Diet rapidly and reproducibly alters the human gut microbiome Nature 505(7484), 559 –563. https://dx.doi.org/10.1038/nature12820

8. Carbonetto, B., Fabbro, M., Sciara, M., Seravalle, A., Mejico, G., Revale, S., Romero, M., Brun, B., Fay, M., Fay, F., Vazquez, M. (2016). Human Microbiota of the Argentine Population-A Pilot Study Frontiers in Microbiology 7(e1500183), 174 –5. https://dx.doi.org/10.3389/fmicb.2016.00051

9. Comeau, A., Douglas, G., Langille, M. (2017). Microbiome Helper: a Custom and Streamlined Workflow for Microbiome Research mSystems 2(1), e00127–16. https://dx.doi.org/10.1128/msystems.00127-16

10. Lozupone, C., Knight, R. (2005). UniFrac: a New Phylogenetic Method for Comparing Microbial Communities Applied and Environmental Microbiology 71(12), 8228–8235. https://dx.doi.org/10.1128/aem.71.12.8228-8235.2005

11. Wong, R., Wu, J., Gloor, G. (2016). Expanding the UniFrac Toolbox PLOS ONE 11(9), e0161196. https://dx.doi.org/10.1371/journal.pone.0161196

12. Kaufman, L. and Rousseeuw, P.J. (1990) Partitioning around Medoids (Program PAM). In: Kaufman, L. and Rousseeuw, P.J., Eds., Finding Groups in Data: An Introduction to Cluster Analysis, John Wiley & Sons, Inc., Hoboken, 68–125.

13. García-Jiménez, Beatriz, and Mark D. Wilkinson. 2018. “Automatic Definition of Robust Microbiome Sub-States in Longitudinal Data.” PeerJ Preprints 6:p e26657v1. doi:10.7287/peerj.preprints.26657v1.

14. Rousseeuw, P. (1987). Silhouettes: A graphical aid to the interpretation and validation of cluster analysis Journal of Computational and Applied Mathematics 20(), 53–65. https://dx.doi.org/10.1016/0377-0427(87)90125-7

15. Tibshirani, R., & Walther, G. (2005). Cluster Validation by Prediction Strength. Journal of Computational and Graphical Statistics, 14(3), 511–528. https://doi.org/10.1198/106186005x59243

16. Real, R., & Vargas, J. M. (1996). The probabilistic basis of Jaccard’s index of similarity. Systematic biology, 45(3), 380–385.

17. Bolyen, E., Rideout, J.R., Dillon, M.R. et al. Reproducible, interactive, scalable and extensible microbiome data science using QIIME 2. Nat Biotechnol 37, 852–857 (2019). https://doi.org/10.1038/s41587-019-0209-9

18. Sonnenburg, E., Sonnenburg, J. (2019). The ancestral and industrialized gut microbiota and implications for human health Nature Reviews Microbiology 17(6), 383–390. https://dx.doi.org/10.1038/s41579-019-0191-8

19. Keohane, D., Ghosh, T., Jeffery, I., Molloy, M., O’Toole, P., Shanahan, F. (2020). Microbiome and health implications for ethnic minorities after enforced lifestyle changes Nature Medicine. 26, 1089–1095 (2020) https://dx.doi.org/10.1038/s41591-020-0963-8

