## Supplementary material for "Generation of a robust reference gut microbiome dataset for an urban population in Argentina optimized by a machine learning approach": Rohr_et_al_suppl

Cristian Rohr[1], Mariela Sciara[2] ,Bianca Brun[1] ,Paula Burdisso[3], Rodolfo Rasia[3,5], Alejandro Vila[3,5], Coral de Val Muñoz[4], Fabian Fay[1,2], Martín P. Vazquez[1,5,*]

[1]Héritas, Rosario, Argentina,
[2] CIBIC, Rosario, Argentina
[3]PLABEM-IBR-CONICET, Rosario, Argentina
[4]University of Granada, Department of Computer Science, Granada, Spain,
[5]CONICET, Argentina.

*Corresponding Author:


Héritas
Medicina de precisión
CIBIC + INDEAR

**HERITAS MICROXPLORA CONTROL DB**

**VOLUNTEERS INFORMATION - Epidemiologic and clinical data**

---

**Date**:

**Name:**

**Adress**

**City:**

**Hábitat:** rural / urban

**Date of birth:**

**Age:**

**Sex:** Female / male

**Nationality:**

**Ethnicity:**

**Weight (kg):**

**Hight (m):**

**BMI ($W/H^2$):**

**Family Relationship with Other Study Volunteers:** yes/ no

**Presence of Twin:** yes/no

**Time of Residence in the Locality:**

**Living under the same roof with others:** yes / no

#### GENERAL BACKGROUND

**Occupation:**

**Smoker:** no / yes - N° cigarettes/day:

**Hours of sleep per day** (on average):

**Dominant hand:** Right / Left

**Cosmetic use:** no /yes - frequency:

**Use of deodorant:** no / yes

**Frequency of dental brushing:**

**Physical activity:** no / yes - frequency and location:

**Swimming:** no / yes - frequency:

**Presence of pets or farm animals in your environment:** no / yes –which

**Trips made in the last 6 months:** no / yes - where

#### DIET

**Diet type:** Varied / Vegetarian / Other – Specify

**Main vegetable ingested:**

**Main carbohydrate ingested:** - % of the diet from carbohydrates

**Source of water consumed:** potable water / Mineral water / Other:

**Alcohol consumption:** No / yes - Frequency:

**Ingestion of multivitamin supplements:** No / yes - which:

**Dietary restrictions:** Lactose / Gluten / Others:

**Dietary change in the last 6 months**: No / yes - which:

**Significant weight modifications in last 3 months:** No / yes -Specify:

#### CLINIC HISTORY

**Diagnosed diseases in the last six months:**

**History of lung disorders:** no / yes - which:

**History of respiratory disorders:** no / yes - which

**Amygdala removal surgery:** no / yes – date:

**History of haematological disorders:** no / yes – which

**History of disorders liver:** no / yes - which

**History of kidney disorders:** no / yes - which

**History of urogenital disorders:** no / yes – which

**History of allergic disorders:** no / yes - which

**History of gastrointestinal disorders**:

Chronic intestinal inflammation, ulcerative colitis, Crohn's disease, Irritable colon syndrome (moderate to severe): no / yes - which

Infectious Gastroenteriris, colitis or gastritis, chronic or persistent diarrhea of unknown origin, recurrent infection with *Clostridium difficile* or untreated *Helicobacter pylori* infection: no / yes - which

Chronic constipation: no / yes

Existence of polyps or gastrointestinal or colonic masses, dysplasia or cancer: no / yes - which

Major surgery of the gastrointestinal tract in the previous 5 years: no / yes

Use of medicines containing bismuth subsalicylate in the last 7 days: no / yes

Appendix removal surgery: No / yes - date

#### GESTATIONAL HISTORY (WOMEN ONLY)

**Current pregnancy:** no / yes

**Currently in lactation period:** no / yes

**Number of full-term pregnancies:**

**Number of abortions:**

**Healthy pregnancies:** no / yes – which disorder

**Delivery type:** Natural / Caesarean

**Type of contraception:** Hormonal / IUD / Condom / Other No If

#### MEDICATION

**Chronic medication:** no /yes – which:

**Circumstantial medication:** no /yes – which:

**Homeopathic medication:** no /yes

Existence of polyps or gastrointestinal or colonic masses, dysplasia or cancer: no / yes - which

Major surgery of the gastrointestinal tract in the previous 5 years: no / yes

Use of medicines containing bismuth subsalicylate in the last 7 days: no / yes

Appendix removal surgery: No / yes - date

#### GESTATIONAL HISTORY (WOMEN ONLY)

**Current pregnancy:** no / yes

**Currently in lactation period:** no / yes

**Number of full-term pregnancies:**

**Number of abortions:**

**Healthy pregnancies:** no / yes – which disorder

**Delivery type:** Natural / Caesarean

**Type of contraception:** Hormonal / IUD / Condom / Other No If

#### MEDICATION

**Chronic medication:** no /yes – which:

**Circumstantial medication:** no /yes – which:

**Homeopathic medication:** no /yes

| Code | Laboratory Clinical Analysis |
|---|---|
| 475 | HEMOGRAM |
| 412 | GLYCEMIA |
| 902 | UREMIA |
| 192 | CREATININE |
| 839 | Na |
| 753 | K |
| 168 | Cl |
| 873 | GOT |
| 874 | GPT |
| 420 | GAMMA GLUTAMIL TRANSPEPTIDASE (GGT) |
| 357 | ALKALINE PHOSPHATASE |
| 110 | TOTAL DIRECT AND INDIRECT BILIRRUBINE |
| 174 | TOTAL CHOLESTEROL |
| 260 | HDL |
| 2601 | LDL |
| 876 | TRIGLICERIDS |
| 503 | HBS AG |
| 495 | ANTI HCV |
| 63 | ANTI HIV |
| 865 | TSH |
| 121 | HCG |

Figure Legend

Clinical lab testing of volunteers. Each point in the continuous line represents a participant.

Yellow lines represent upper threshold limits and red lines lower threshold limits. The most relevant measures were included in the plots.

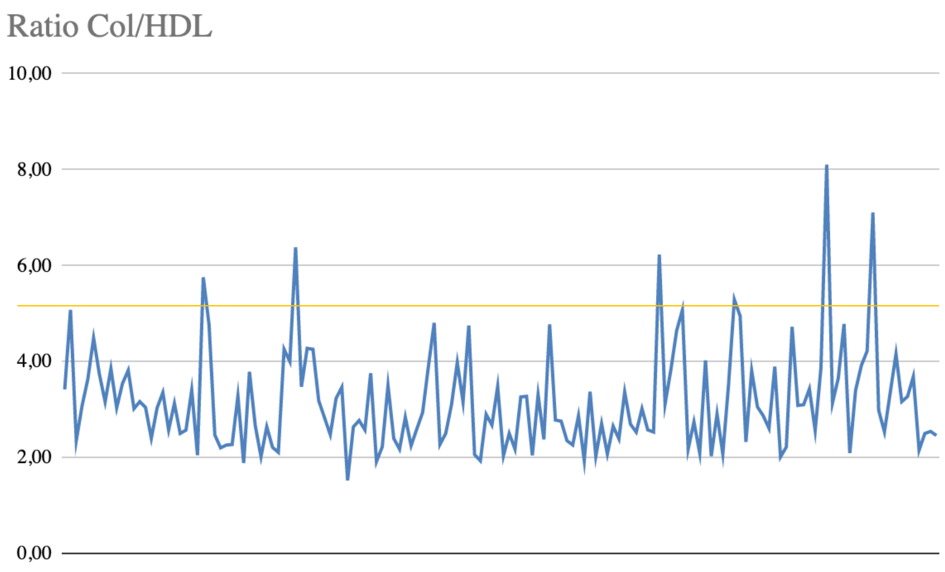


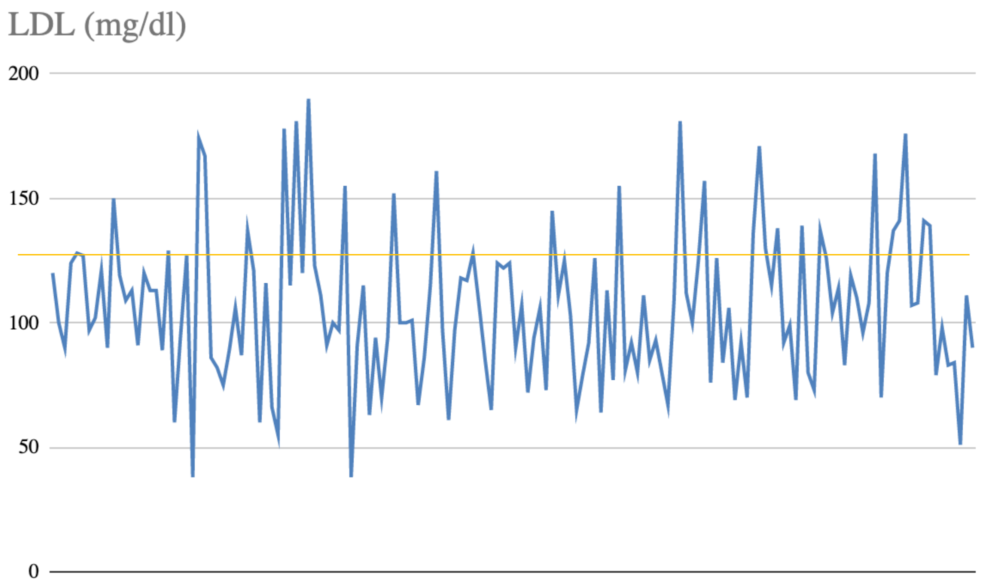


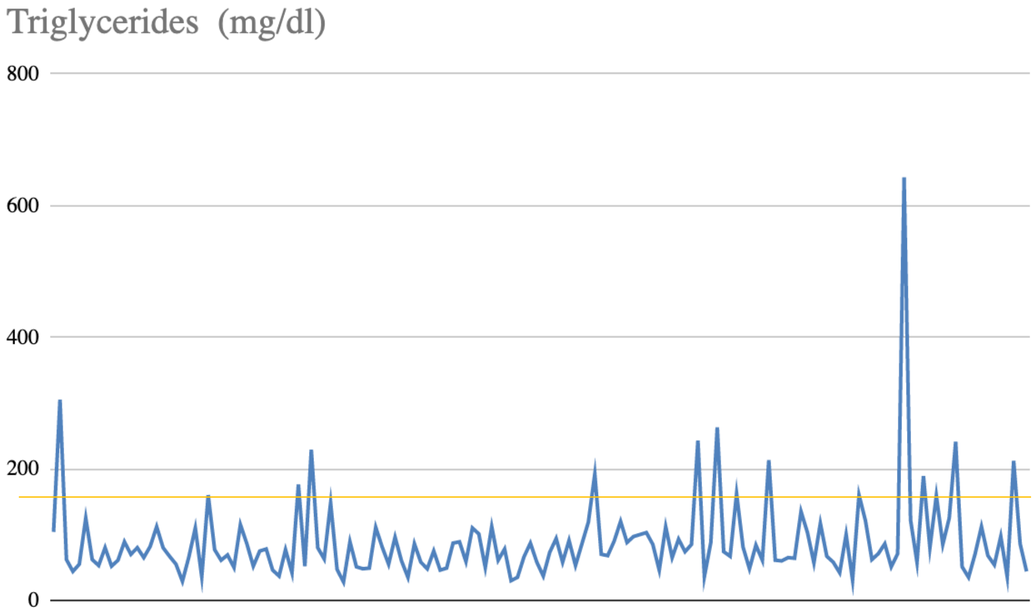


Rohr et al.

### Lipidic levels

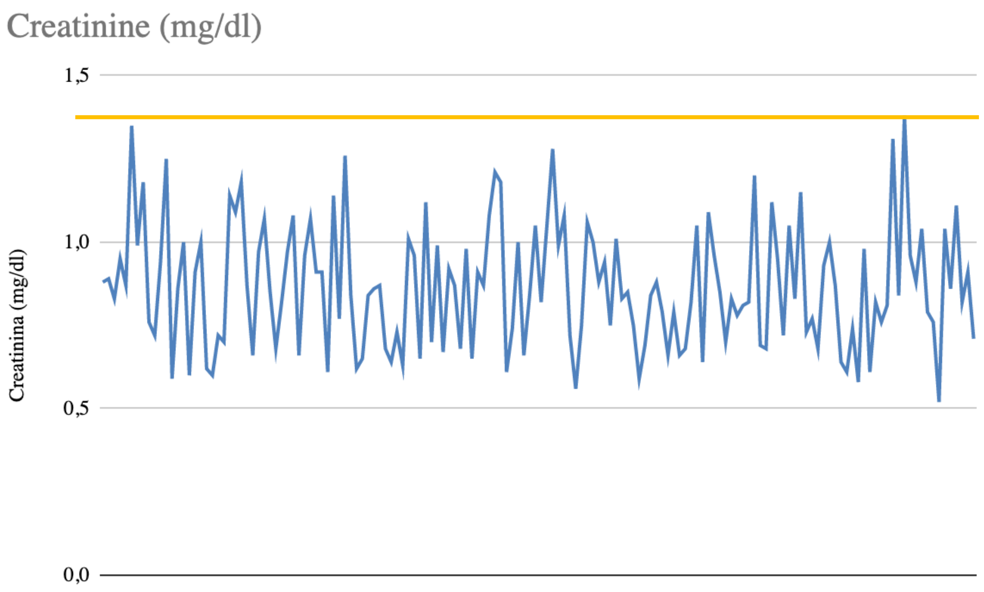
Creatinine (mg/dl)
1,5
1,0
0,5
0,0
Creatinina (mg/dl)

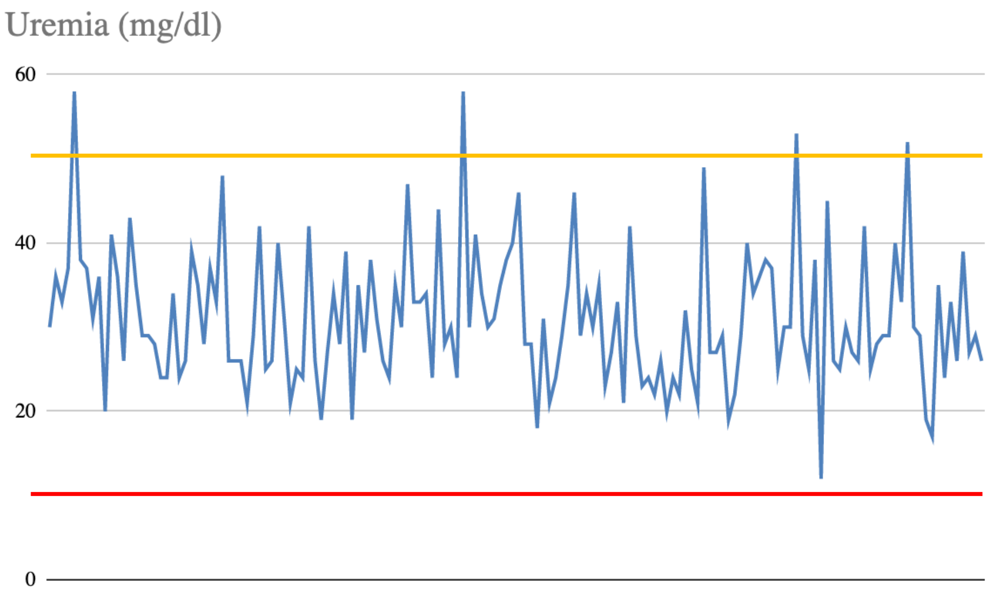
Uremia (mg/dl)
60
40
20
0

Rohr et al.

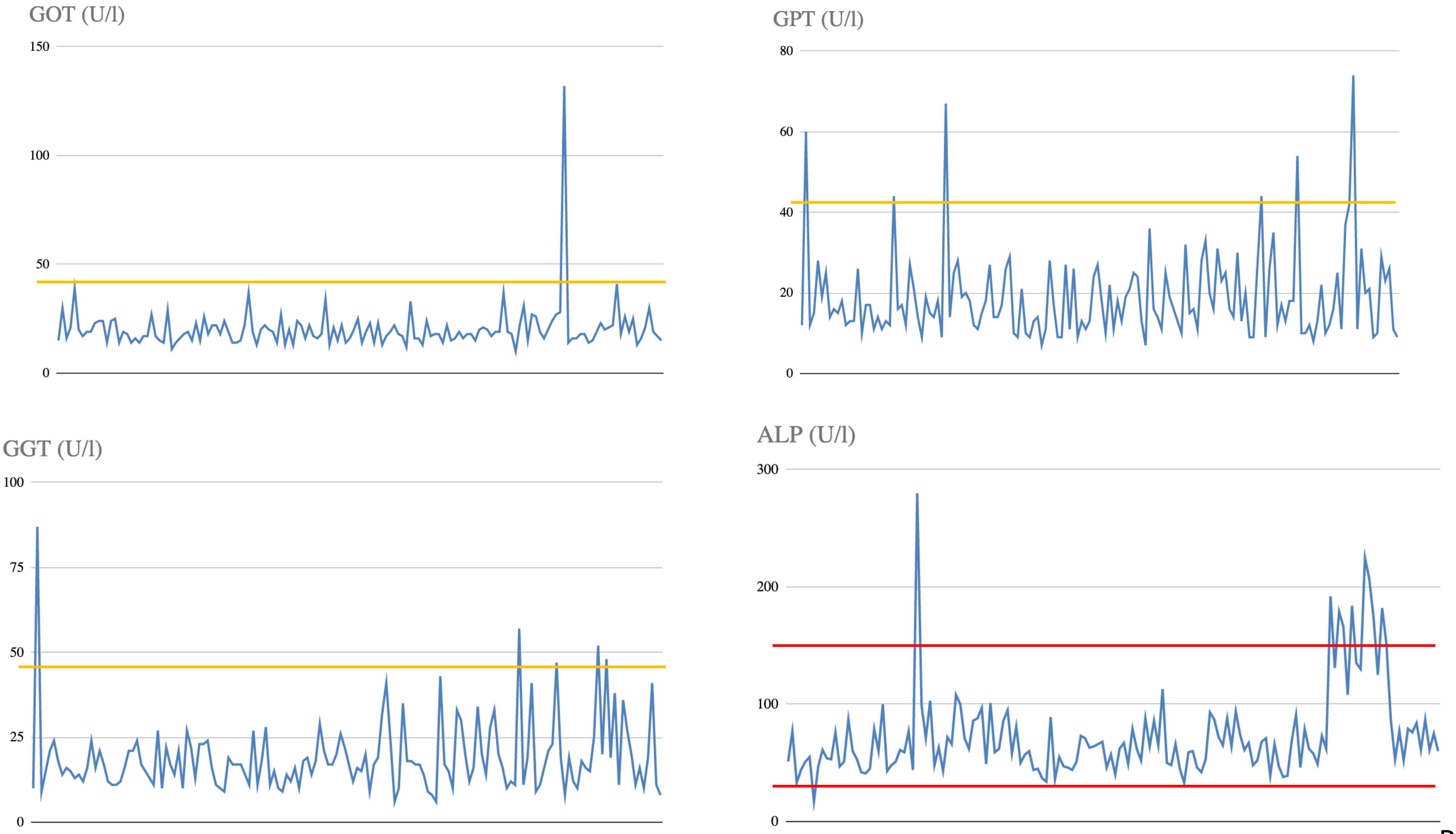
GOT (U/l)
150
100
50
0
GPT (U/l)
80
60
40
20
0
GGT (U/l)
100
75
50
25
0
ALP (U/l)
300
200
100
0

Rohr et al.

#### MicroXplora Argentine Reference Dataset (MARD)
#### 172 volunteer samples

### **Plasma Metabolome**:
### NMR CPMG spectra

Rohr et al.

# S3

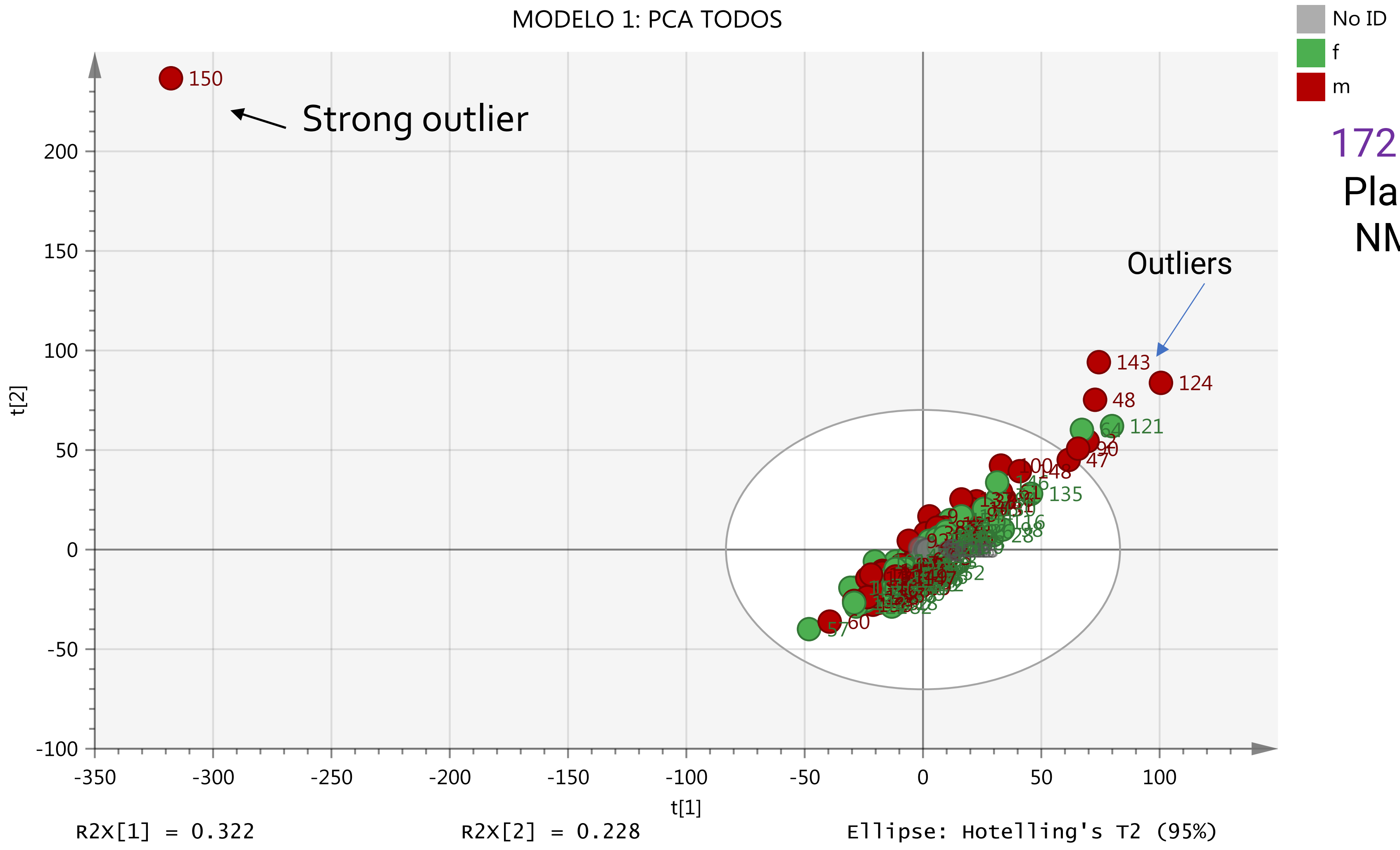


172 volunteer samples

Plasma Metabolome:
NMR CPMG spectra

Rohr et al.

# S4

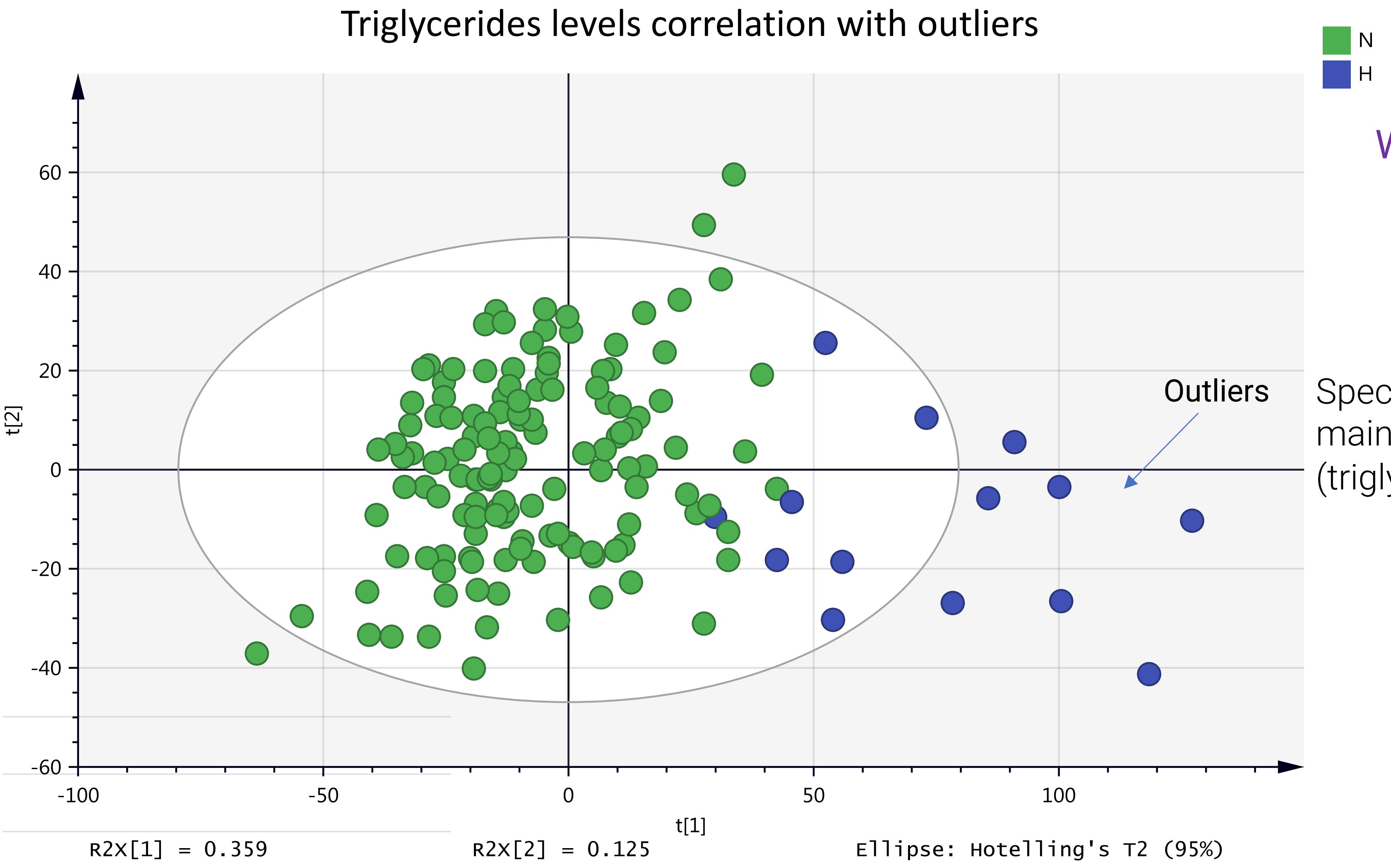


Without strong outlier 171 samples

Plasma Metabolome:
NMR CPMG spectra

Spectra
mainly VLDL
(triglycerides transport)

Rohr et al.

### MicroXplora Argentine Reference Dataset (MARD)
### 172 volunteer samples

**Urine Metabolome**:
NMR NOESY spectra

Rohr et al.

# S5

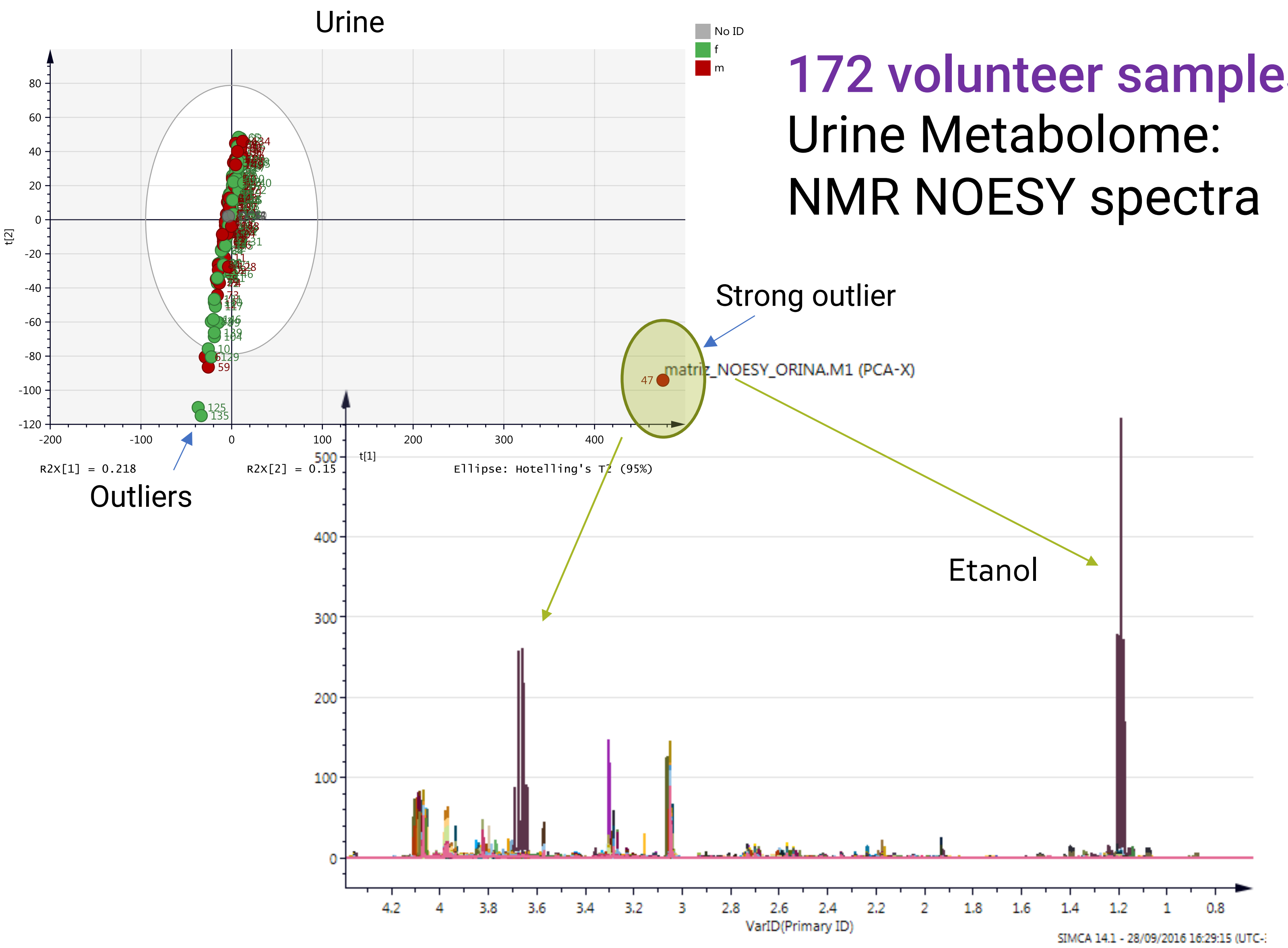


172 volunteer samples

Urine Metabolome:

NMR NOESY spectra

Rohr et al.

S6

# PC1 vs PC2

#### Without the strong outlier (171 samples)

#### Urine Metabolome: NMR NOESY spectra

**Hippuric acid** is the most frequently used biomarker in the biological monitoring of occupational exposure to toluene

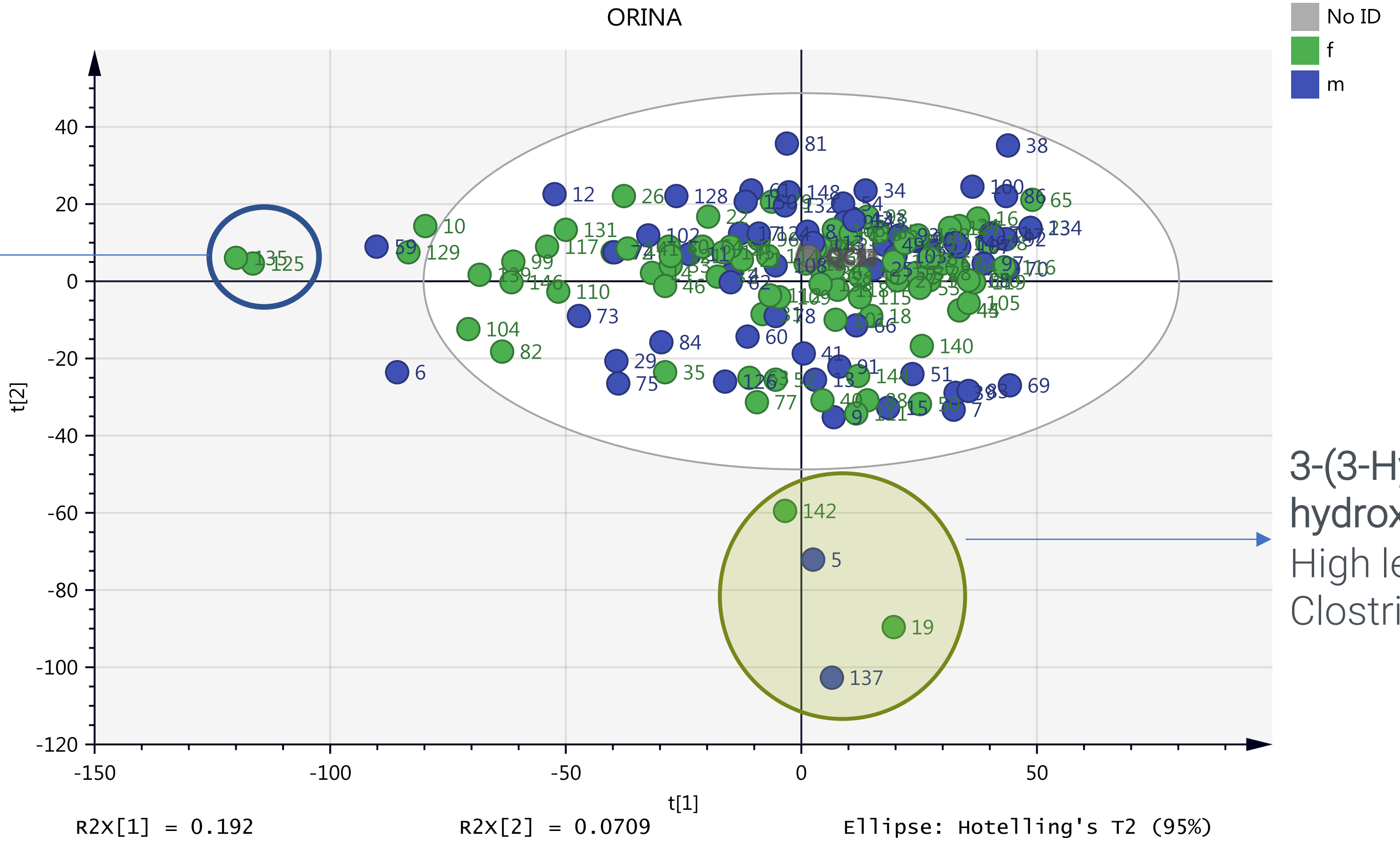


3-(3-Hydroxyphenyl)-3-hydroxypropanoic acid. High levels produced by Clostridia sp in the gut

Rohr et al.

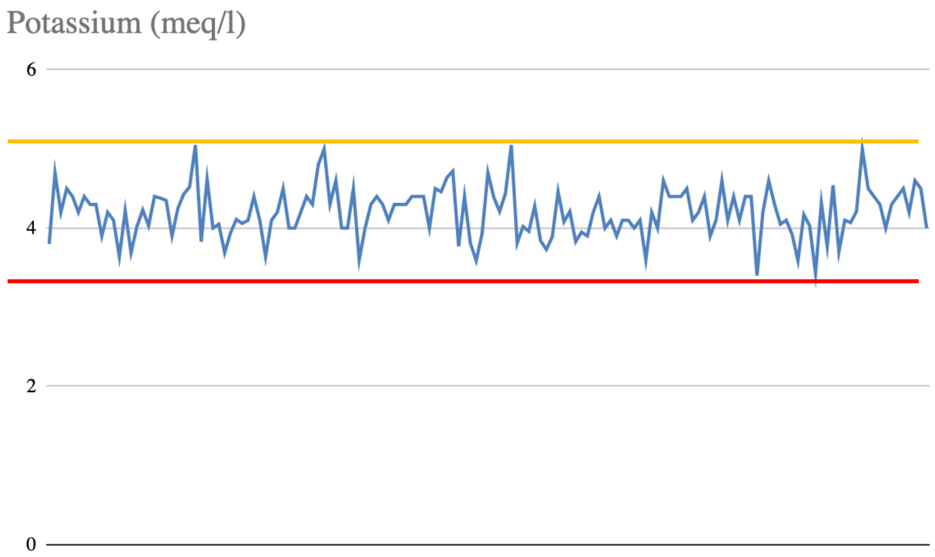


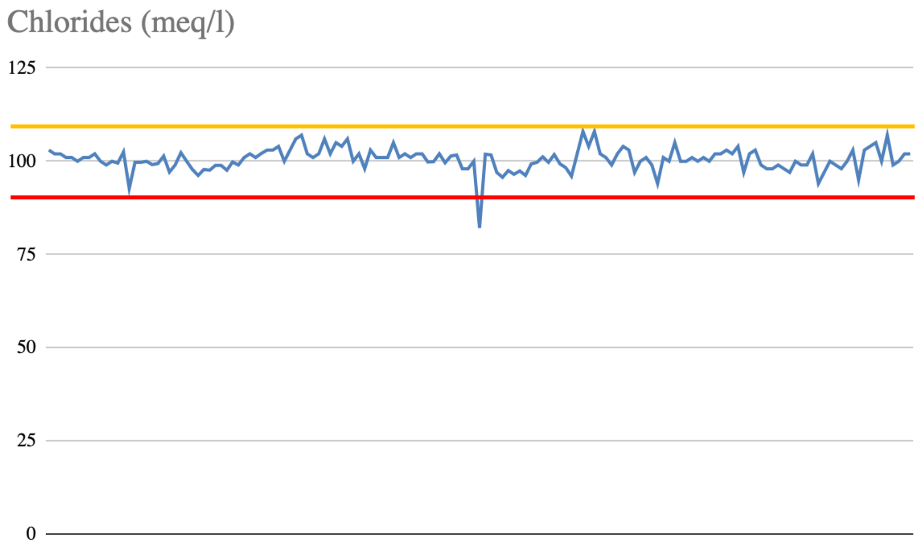


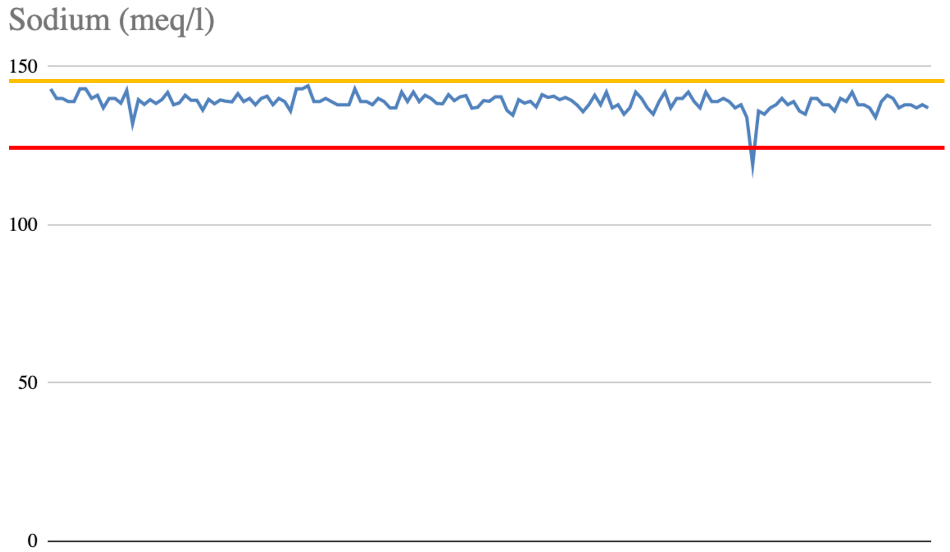


Rohr et al.

#### Electrolyte levels

# S7

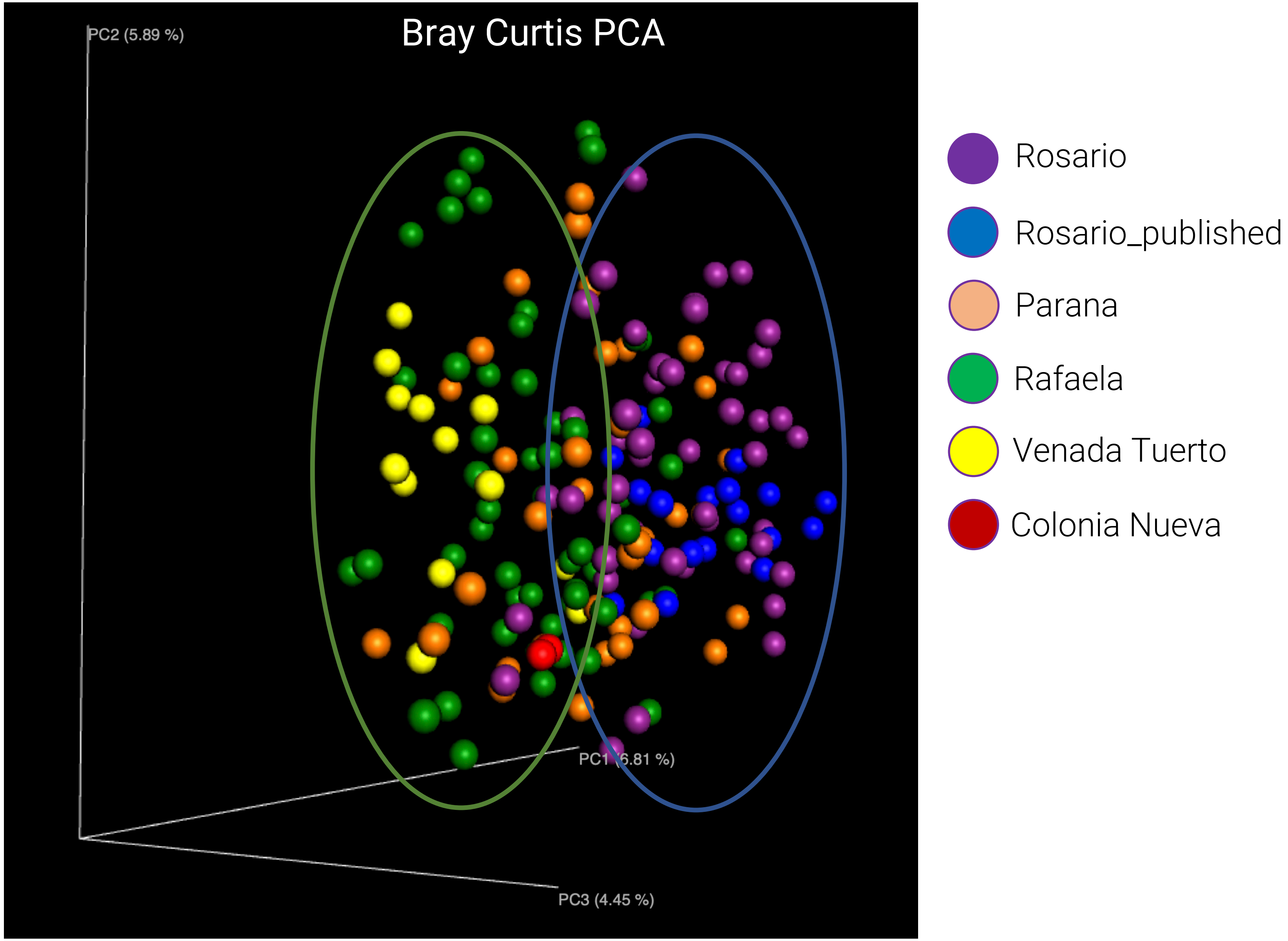
Bray Curtis PCA
PC2 (5.89 %)
PC1 (6.81 %)
PC3 (4.45 %)
Rosario
Rosario_published
Parana
Rafaela
Venada Tuerto
Colonia Nueva

Principal components analysis (PCoA) using Bray-Curtis distance matrix of the gut microbiome data obtained from the four cities. Blue circle labeled Rosario_published corresponds to the gut microbiome from Rosario city previously published by our group using the same methodology (8). Red circle labeled Colonia Nueva refers to 2 volunteers from a rural town near Rafaela.

S8

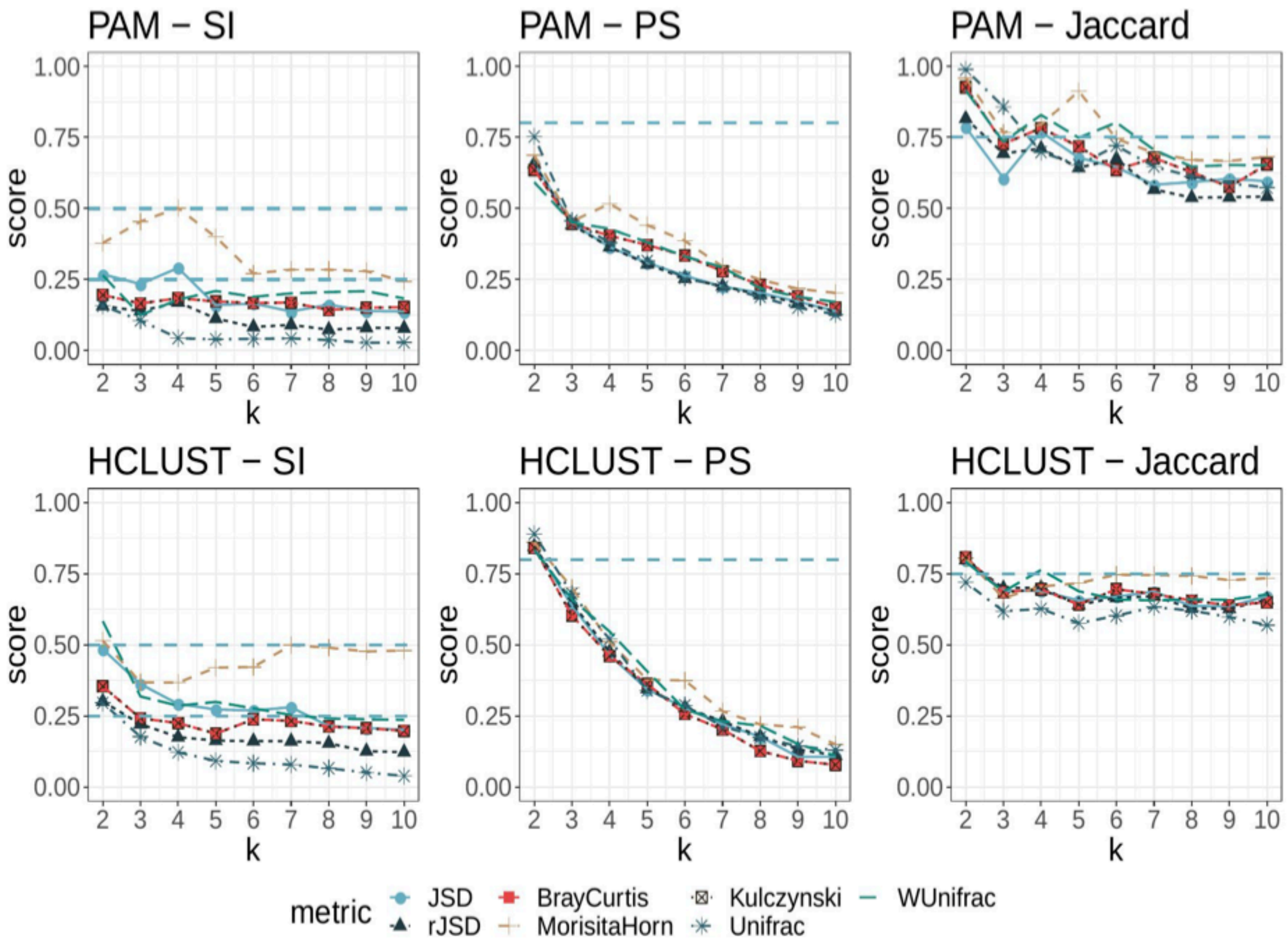
PAM – SI
PAM – PS
PAM – Jaccard
HCLUST – SI
HCLUST – PS
HCLUST – Jaccard
score
k
1.00
0.75
0.50
0.25
0.00
2 3 4 5 6 7 8 9 10
metric
JSD
rJSD
BrayCurtis
MorisitaHorn
Kulczynski
Unifrac
WUnifrac

S9

### Supervised classification models | Results

Results supervised classification models using Random Forests and Support Vector Machines. Alg=Algorithm, Ov=Oversample, Opt=Optimized

| Alg. | Ov. | Opt. | Parameters | Accuracy | MicroF1 | MacroF1 |
|---|---|---|---|---|---|---|
| RF | No | No | mtry = 8; ntree = 1000 | 0.9091 | 0.9524 | 0.8864 |
| RF | Yes | No | mtry = 8; ntree = 1000 | 0.9773 | 0.9885 | 0.9519 |
| RF | No | Yes | mtry = 15; ntree = 5000 | 0.9318 | 0.9647 | 0.9043 |
| RF | Yes | Yes | mtry = 8; ntree = 1000 | 0.9773 | 0.9885 | 0.9519 |
| SVM | No | No | kernel = radial; sigma = 0.0119; C = 1 | 0.7955 | 0.8861 | 0.7681 |
| SVM | Yes | No | kernel = radial; sigma = 0.0278; C = 1 | 0.9318 | 0.9647 | 0.8705 |
| SVM | No | Yes | kernel = poly; degree = 2; scale = 0.001; C = 10 | 0.8864 | 0.9398 | 0.8326 |
| SVM | Yes | Yes | kernel = radial; sigma = 0.001; C = 10 | 0.9545 | 0.9767 | 0.9356 |

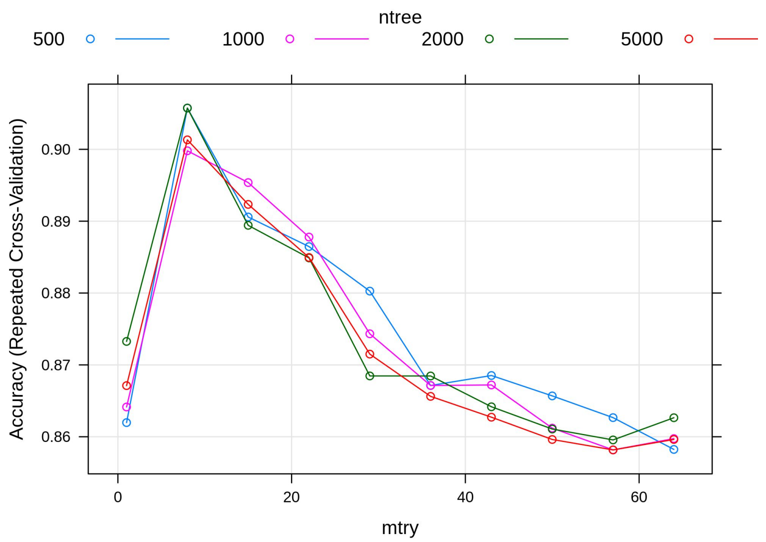


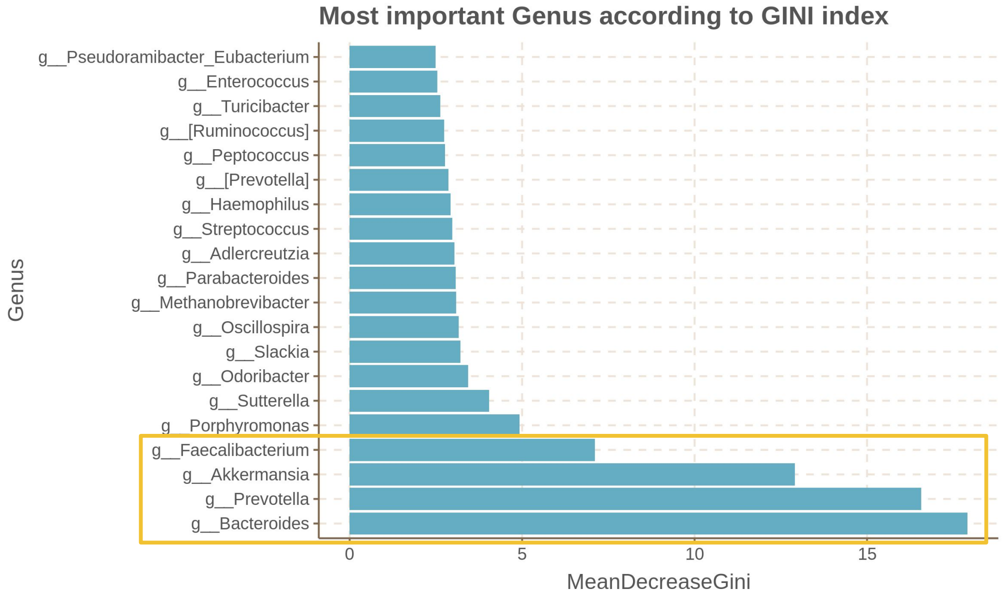

S10

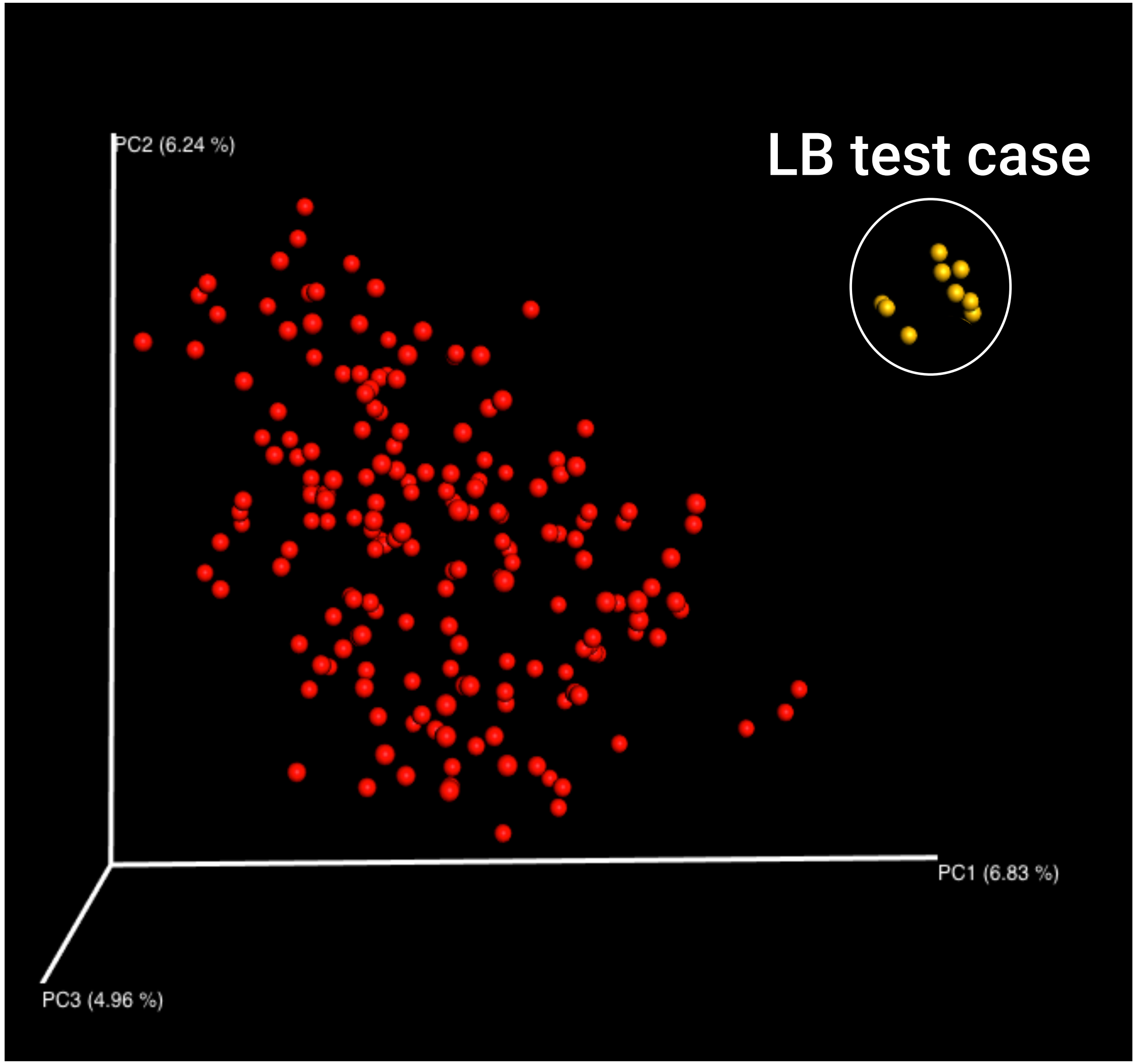


Rohr et al.

### Figure Legend

Bray Curtis distance matrix analysis of the gut microbiome reference datatset (red dots) and the LB test case (yellow dots). LB samples were taken in triplicates at days 1, 7 and 15.
